# The mitochondrial NAD transporter SLC25A51 is a modulator of beta cell senescence and type 2 diabetes

**DOI:** 10.1101/2025.04.09.647991

**Authors:** Byung Soo Kong, Sehi L’Yi, Erika Gabriela Ramirez-Hernandez, Serin Hong, Rosa Weidenspointner, Suyeon Song, Chase Caserta, Young Min Cho, Nora Kory

**Affiliations:** Department of Molecular Metabolism, Harvard T.H. Chan School of Public Health, Boston, MA, USA; Dana-Farber Cancer Institute, Boston, MA, USA; Broad Institute of MIT and Harvard, Cambridge, MA, USA; Department of Biomedical Informatics, Harvard Medical School, Harvard University, Boston, MA 02115; Department of Internal Medicine, Seoul National University College of Medicine, Seoul, Korea; Division of Endocrinology and Metabolism, Department of Internal Medicine, Seoul National University Hospital, Seoul, Korea

## Abstract

Nicotinamide adenine dinucleotide (NAD^+^) is an essential redox cofactor and signaling molecule linked to age-dependent metabolic decline, with its compartmentalization regulated by the mitochondrial carrier SLC25A51. The mechanisms contributing to declining NAD^+^ levels during aging and the consequences of altered NAD^+^ homeostasis across tissues are poorly understood. Here, we show that SLC25A51 is upregulated in aging and aging-associated conditions, particularly in senescent cells. In a mouse model of beta-cell senescence, upregulated SLC25A51 was associated with beta-cell identity loss, senescence progression, and a reduced NAD^+^/NADH ratio. SLC25A51 was elevated following p16^INK4a^-, replicative-, irradiation-, and H_2_O_2_-induced senescence, with NRF2 implicated as a potential transcriptional regulator. Overexpression of SLC25A51, but not a transport-dead mutant, induced senescence factors, while its deletion prevented this effect. Beta-cell-specific deletion of SLC25A51 lowered p16^INK4a^ levels in pancreatic islets, circulating insulin, and glucose levels, improving insulin sensitivity and indicating its role in cellular senescence and the metabolic control of beta-cell function.

## Introduction

Aging is a complex phenomenon marked by changes in biological processes and metabolism, increasing disease susceptibility^1^. The redox cofactor nicotinamide adenine dinucleotide (NAD^+^) links metabolic control with cellular and epigenetic states. Disrupted NAD^+^ levels or redox balance are observed in aging and age-related diseases across model organisms and tissues^2–15^ (reviewed in ^16^). Altered NAD^+^ levels and NAD^+^/NADH redox balance are also linked to mitochondrial dysfunction and cellular senescence, which are key hallmarks of aging (reviewed in ^17^). Senescent cells are characterized by irreversible cell cycle arrest in response to aging-related stress^5,18–29^. While it is well-established that NAD^+^ levels decline during aging and in age-related diseases^2,3,5,6,14,30–35^, the processes contributing to this decline across different tissues, as well as the effects of altered NAD^+^ levels on cell function, metabolism, and state, remain incompletely understood.

Likewise, the phenotype of senescent cells varies depending on the physiological context. Senescence protects against tumorigenesis by stably arresting mitotic cells, such as fibroblasts or immune cells, but it also contributes to changes in metabolic activities and gene expression associated with inflammation and aging^36^. Postmitotic cells, which are unreplaceable or infrequently replaced with age, including neurons, pancreatic beta cells and oocytes, are particularly vulnerable to metabolic stress and can also become senescent^37,38^. Pancreatic beta-cell senescence is of particular interest as it is increasingly recognized as a contributor to the pathogenesis of both type 1 (T1D) and type 2 diabetes (T2D)^19,27,39–41^. This has highlighted the targeting of senescent beta cells as a promising therapeutic approach for diabetes^42^. Despite these advancements, major questions remain regarding the molecular mechanisms underlying beta-cell senescence, the identification of specific markers differentiating senescent from non-senescent beta-cell subsets, and the translation of proof-of-concept therapies into novel diabetes treatments.

Cellular NAD^+^ levels and NAD^+^/NADH ratios are altered in beta-cell aging and diabetes^4,43^. Cercillieux et al. demonstrated that whole-body knockout of nicotinamide riboside kinase 1 (Nrk1) in mice reduced pancreatic NAD^+^ levels, impaired glucose tolerance, and reduced insulin secretion and sensitivity upon a high-fat diet^4^. Similarly, Murao et al. found that nicotinamide mononucleotide adenylyl transferase 2 (Nmnat2), a cytosolic NAD^+^ synthesis enzyme, was induced in pancreatic beta cells in aged and diabetic mouse models, possibly as an adaptive response to increased glycolytic flux, compensating for decreased NADH re-oxidation^43^. Conversely, NMNAT2 suppression improved beta-cell identity and function under increased glycolytic flux, potentially by altering NAD^+^ availability for nuclear NAD^+^ synthesis and its dependent enzymes, Sirt1 and PARP1, in a mechanism analogous to changes in compartmentalized NAD^+^ synthesis during adipogenesis^44^. Overall, these findings highlight the critical role of NAD^+^ homeostasis and compartmentalization in regulating beta-cell function and the diabetes pathogenesis. However, the mechanisms by which compartmentalized NAD^+^ pools are established within cells and how they affect NAD^+^- dependent processes in relation to aging, cell state, function, and metabolism remain unclear. Recently, we and others identified SLC25A51 as the mitochondrial NAD^+^ transporter in metazoans, which plays a critical role in regulating mitochondrial NAD^+^ levels and intracellular NAD^+^ compartmentalization. Through its role in shifting cellular NAD^+^ compartmentalization, SLC25A51 has also been shown to alter NAD^+^-dependent metabolic pathways, DNA repair, and cancer cell proliferation by uncoupling the mitochondrial NAD^+^/NADH ratio^45–47^.

Given the complex interplay between NAD^+^ and the control of cell state and aging, we investigated how the expression of SLC25A51 changes during aging across tissues and under various conditions in metazoans. Using single-cell RNA sequencing (scRNA-seq) of pancreatic islets in a T2D mouse model, we found that SLC25A51 is upregulated in the early to mid-stages of senescent beta cells and modulates NAD^+^ compartmentalization, beta-cell state, and beta-cell function. Furthermore, the beta-cell-specific deletion of SLC25A51 improved insulin sensitivity and glucose homeostasis in a T2D model, demonstrating SLC25A51’s potential as a therapeutic target for diabetes treatment.

## Results

### Induction of mitochondrial NAD^+^ transporters is an evolutionarily conserved feature of chronological aging

To investigate the link between NAD^+^ compartmentalization, mitochondrial function, metabolic regulation, and aging, we analyzed the expression of mitochondrial NAD^+^ transporters using publicly available datasets from various model organisms undergoing chronological aging, as well as cells undergoing senescence *in vitro*. Mitochondrial NAD^+^ transport is mediated by *Ndt1* and *Ndt2* in yeast and plants^48,49^. Recently, we and others identified SLC25A51 as the mitochondrial NAD^+^ transporter in mammals^50–52^. We found that expression of the SLC25A51 ortholog, *Ndt2*, was upregulated in *Arabidopsis thaliana* and *Saccharomyces cerevisiae* during chronological aging (**Figure 1A**), in line with a previous study^53^. Furthermore, SLC25A51 was upregulated in some human cell types undergoing replicative, oxidative stress-induced, or DNA damage-induced senescence such as lung fibroblasts (WI38^54^, MRC5^55^, IMR90^56^), eye epithelial (RPE-1^57^), liposarcoma (LS8817^58^), and bone marrow cells (MSC^56^), but not in endothelial HUVEC, dermal fibroblast (HDF161^59^), and foreskin fibroblast (HFF^55^) (**Figure S1B**). To validate these findings and identify tissues and cell types where SLC25A51 accumulates over time in mammals, we examined SLC25A51 protein levels in various isolated organs and tissues from 90-week-old male and female mice. SLC25A51 was ubiquitously expressed, with the highest mRNA levels observed in the testis, breast, pancreas, and adipose tissue (**Figure S1A**). In aged mice, we detected SLC25A51 protein in pancreatic islets and the liver, but not in the gonads, kidney, spleen, or adipose tissue (**Figure 1B**). Notably, SLC25A51 protein levels increased in pancreatic islets with age, particularly in male mice, whereas other mitochondrial proteins did not show a similar trend (**Figures 1B-1D**). Mining a published scRNA-seq dataset from humans, we found that SLC25A51 was predominantly expressed in beta cells, with minimal expression in other pancreatic islet cell types such as alpha cells, delta cells, or pancreatic polypeptide cells (**Figure S1B**). Immunofluorescence (IF) staining of pancreatic tissues from aged mice confirmed that SLC25A51 accumulated in beta cells, the predominant cell type in mouse islets marked by insulin expression, but not in glucagon-producing alpha cells (**Figures 1E, 1F**). In line with the increased prevalence of T2D and beta-cell senescence with aging^19,39^, SLC25A51 accumulation in 90-week-old mice was accompanied by increased pancreatic senescence-associated beta-galactosidase (SA-beta-Gal) activity, p16^INK4A^ and p21^Cip^^1/Waf1^ levels (markers of senescent beta cells^19,20,22,39,60^), body mass, random blood glucose levels, and serum senescence-associated secretory phenotype (SASP) factors, such as CXCL10 (**Figures 1G, H-J, S1F**).

**Figure 1.**
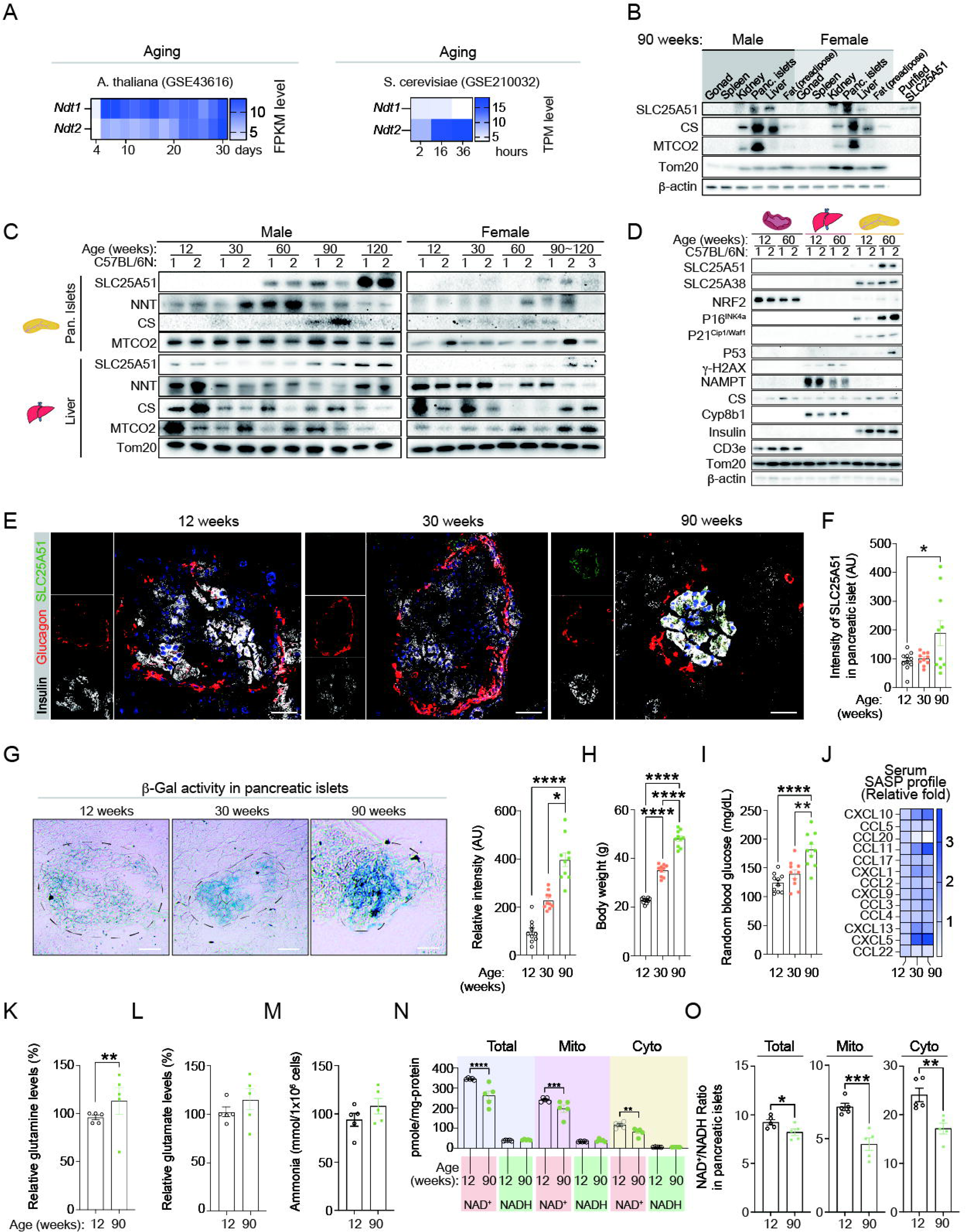
Induction of mitochondrial NAD^+^ transporters is an evolutionarily conserved feature of chronological aging. (**A**) Published datasets were analyzed for mRNA levels of SLC25A51 orthologs (*Ndt1*, *Ndt2*) in *A. thaliana* (GSE43616) and *S. cerevisiae* (GSE210032). (**B**) Levels of SLC25A51 and other mitochondrial proteins, as well as β-actin, were analyzed in various tissues harvested from 90-week-old C57BL/6N male and female mice (gonad, spleen, kidney, pancreatic islet cells, liver, and fat [preadipocytes]). € Levels of SLC25A51 and other mitochondrial proteins were analyzed in pancreatic islets and livers from 12-, 30-, 60-, 90-, and 120-week-old C57BL/6N male and female mice (n=2∼3/group). (**D**) Levels of SLC25A51 protein, senescence markers (p16^INK4a^, P21^cip1/waf1^, p53, γ-H2AX, NAMPT), mitochondrial markers (Tom20, Citrate synthase [CS]), Cyp8b1 (hepatocyte marker), insulin (beta-cell marker), and CD3e (T-cell marker) were analyzed in tissue-derived primary cells (splenocytes, hepatocytes, pancreatic islet cells) isolated from 12- and 60-week-old male C57BL/6N mice (n=2/group). (**E-J**) Twelve-, 30-, and 90-week-old C57BL/6N male mice (n=10/group) were analyzed for (**E, F**) IF staining of SLC25A51, insulin, glucagon, and DAPI; Scale bars, 50 μm. (**G**) Beta-Gal activity in pancreas tissue sections; Scale bars, 50 μm. (**I**) Body weight, (**J**) random blood glucose levels, and (**K**) serum SASP profile. One-way ANOVA; error bars are SEM. *p < 0.05, **p < 0.01, ****p < 0.0001. (**K-O**) Pancreatic islet cells were isolated from 12- and 90-week-old C57BL/6N mice (n=5/group) to measure (**K**) glutamine, (**L**) glutamate, (**M**) cellular ammonia, and (**N, O**) NAD^+^ and NADH levels and NAD^+^/NADH ratio. Two-tailed *t*-test; error bars are SEM. *p < 0.05, **p< 0.01, ***p < 0.001.

### SLC25A51 accumulation in beta cells contrasts with a decline in mitochondrial NAD^+^ during aging in mice

At the whole-tissue level, NAD^+^ levels decline with aging in the pancreas and other organs^16^. Given the increased expression of the mitochondrial NAD^+^ transporter SLC25A51 in pancreatic beta cells with aging, we assessed whether mitochondrial NAD^+^ levels in beta cells changed concomitantly. We found that pancreatic islet cells isolated from 12-week, 30-week, and 90-week-old mice consisted of 80% insulin-producing beta cells, consistent with previous reports^61,62^ (**Figure S1C**). Compared to 12-week-old mice, 90-week-old mice exhibited a decline in total NAD^+^ levels, as previously reported^4,43^, as well as the NAD^+^/NADH ratio, in whole-cell lysates (total), cytosolic fractions, and mitochondrial fractions (**Figures 1N, 1O**). Notably, this decline occurred despite the increased expression of SLC25A51. We also observed other metabolic features characteristic of senescent cells and tissues, including increased glutamine, glutamate, and cellular ammonia levels in whole-cell lysates, indicative of enhanced glutaminolysis^63^ (**Figures 1K-1M**). These findings indicate that both whole-tissue and mitochondrial NAD^+^ levels decrease in pancreatic islets during chronological aging, in contrast to the increase in SLC25A51 expression.

### Pancreatic islet cells isolated from S961-treated C57BL/6N mice are transcriptionally heterogeneous and exhibit increased senescent beta-cell features marked by mitochondrial inner membrane transporter gene signatures

We hypothesized that SLC25A51 accumulates in beta cells as they become senescent, contributing to metabolic changes and the senescence phenotype. To test this hypothesis and investigate SLC25A51 expression and function, we performed scRNA-seq using a previously established mouse model of T2D and acute beta-cell senescence induced by the insulin receptor antagonist S961^19,39,40,64^. We administered systemic treatment with S961 or PBS to C57BL/6N mice at 28 weeks of age (n=10/group) using an osmotic pump for two weeks (**Figure 2A**). S961 treatment induced hyperglycemia (12-week-old control [12W-C]: 111.20±12.04 mg/dL; 30-week-old control [30W-C]: 104.20±14.69 mg/dL; 30-week-old S961 treated [30W-S]: 229.70±98.21 mg/dL) and hyperinsulinemia (12W-C: 6.30±0.04 ng/ml; 30W-C: 15.91±0.19 ng/ml; 30W-S: 25.80±1.39 ng/ml) compared to both 12- and 30-week-old control mice (**Figures 2B-2F**). Overall, diabetes incidence was increased in S961-treated mice compared to control mice at 30 weeks (70% versus 0%, n=10/group) (**Figure 2D**). These effects were independent of body weight (**Figure 2C**).

**Figure 2.**
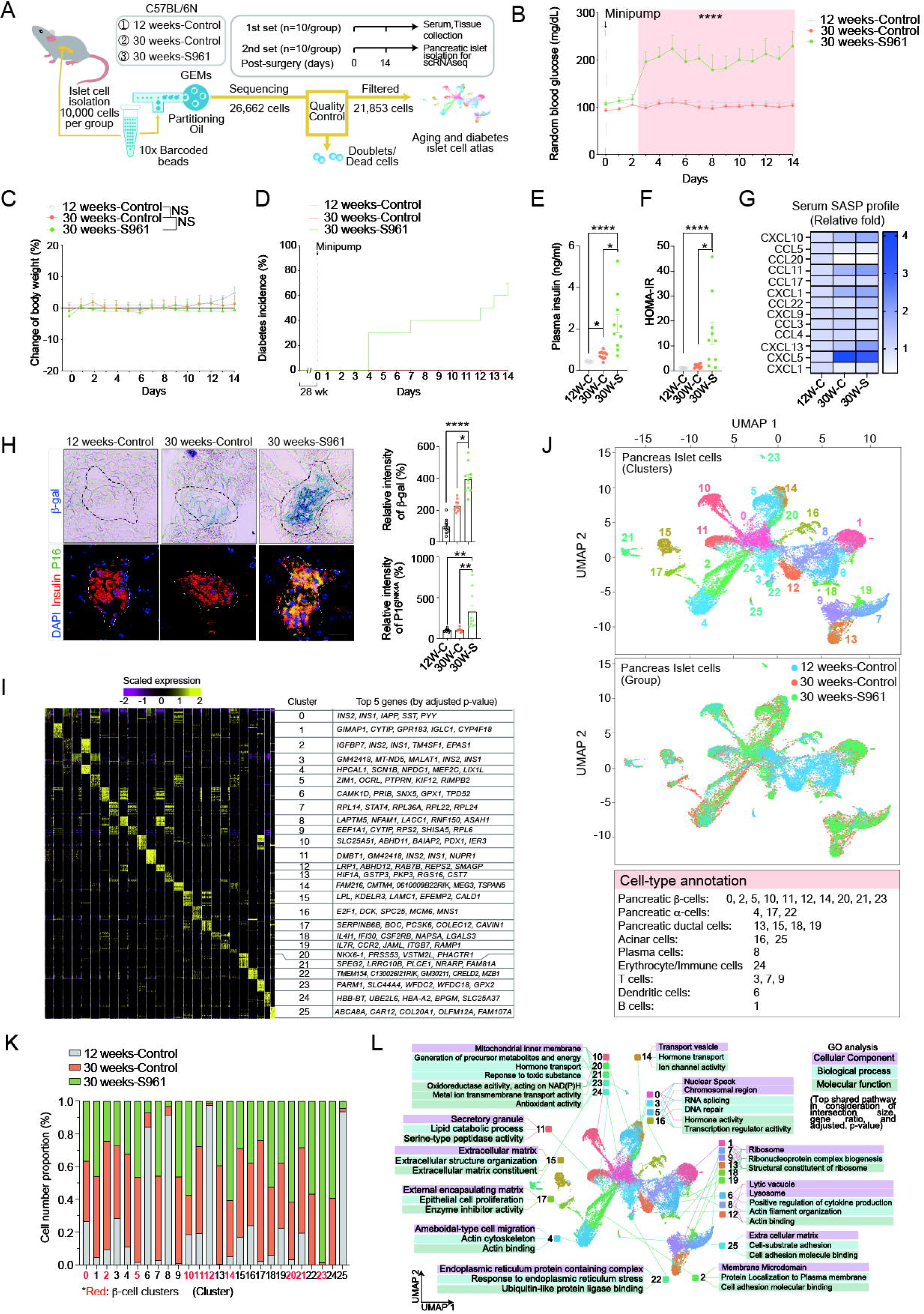
Pancreatic islet cells isolated from S961-treated C57BL/6N mice are transcriptionally heterogeneous, and exhibit increased senescent beta-cell features marked by mitochondrial inner membrane transporter gene signatures. Twelve-week-old C57BL/6N mice were bred and fed *ad libitum* until 28 weeks of age. (**A**) At 28 weeks, the mice were implanted with a mini-pump containing either vehicle or S961 (n=10). Mice were monitored for 2 weeks to analyze (**B**) random blood glucose levels and (**C**) body weight; two-way repeated measures ANOVA; error bars represent SEM. ****p < 0.0001. Additionally, (**D**) diabetes incidenc€(**E**) plasma insulin levels, (**F**) HOMA-IR, and (**G**) serum SASP profiles (n=10/group) were assessed; one-way ANOVA; error bars are SEM. *p < 0.05, ****p < 0.0001. After 2 weeks of monitoring, 12-week-old control (12W-C), 30-week-old control (30W-C), and 30-week-old S961-treated (30W-S) mice were sacrificed (n=10/group). Subsequently, beta-Gal, p16^INK4A^, and insulin staining were performed in pancreatic islets on (**H**) frozen sections and (**Figure S1D**) live cells using flow cytometry to confirm S961-induced β-cell senescence. Scale bars, 50 μm. One-way ANOVA; error bars are SEM. *p < 0.05, **p < 0.01, ****p < 0.0001. Using a second set of animals (n=10/group), we conducted (**A, H, I, J**) pancreatic islet scRNA-seq. From this experiment, 21,853 qualified pancreatic islet cells were obtained from three groups. The data were used to define (**J**) the beta-cell senescence-induced diabetes islet cell atlas and (**I**) the top five differentially expressed genes. An overview of the beta-cell senescence-induced diabetes islet cell atlas is presented, showing clusters (upper panel), group distributions (middle panel), and cluster annotation (lower panel). Cell-type annotation in scRNA-seq suggests clusters 0, 2, 5, 10, 11, 12, 14, 20, 21, and 23 represent pancreatic beta-cells. (**K, L**) Analyses in all clusters in three groups (12W-C, 30W-C, and 30W-S) were performed. (**K**) Cell number and proportion (%) were evaluated in each cluster and group (**See also Figure S1E**). (**L**) GO (GO:CC, GO:BP, GO:MF) analyses in scRNA-seq data showed that 26 different clusters can be categorized into 12 different groups based on GO:CC.

To assess beta-cell senescence, we examined SASP production, SA-beta-Gal activity, and p16^INK4A^ levels in pancreatic beta cells. SASP-related chemokines, such as CXCL10, CCL4, CCL3, CXCL1, and CCL5, have previously been reported to increase in hydrogen peroxide (H_2_O_2_)-induced senescent MIN6 beta cells^19^. Furthermore, human and mouse senescent beta cells secrete SASP factors such as eNAMPT, CCL2, CCL4, CXCL2, and CXCL3^40^. In addition, CCL20, CCL11, CCL17, CCL22, CXCL9, CXCL13, and CXCL5 are critical SASP factors in different tissue types and disease models^65–69^, although they have not been evaluated in S961-treated diabetic mouse models. We observed that CCL2 was elevated in the 30W-S group compared to both the 12W-C (p=0.0008) and 30W-C (p=0.0165) groups, while CXCL10 was slightly elevated, reaching significance only compared to 12-week-old control animals (12W-C: p<0.0001; 30W-C: p=0.0651) (**Figures 2G, S1G**). Other SASP factors did not show significant differences between the 30W-C and 30W-S groups (**Figures 2G, S1G**). Staining for CD45, a pan-lymphocyte marker, and SA-beta-Gal activity in pancreatic islet cells revealed a 2- and 4-fold increase in CD45^Negative^ SA-beta-Gal^positive^ beta cells in the 30W-S group compared to the 30W-C and 12W-C groups, respectively (**Figure S1D**). Staining of pancreatic tissue confirmed increased SA-beta-Gal activity and p16^INK4A^ expression in the 30W-S group in insulin^positive^ pancreatic beta cells (**Figure 2H**). In summary, these data indicate that systemic S961 treatment in C57BL/6N mice, as previously described for C57BL/6J mice^19,39,64^, induces beta-cell senescence by increasing *Igf1r* and *p16^Ink4a^* mRNA levels and insulin levels, contributing to the development of insulin resistance and T2D.

To specifically identify and profile senescent beta-cell subsets, we generated single-cell atlases for pancreatic islets isolated from three groups: 12W-C, 30W-C, and 30W-S mice (**Figures 2A, 2I, 2J**). To define each cell type, we processed the sequencing data using the Seurat R package for quality control, normalization, batch effect correction, and clustering. Each cell type was annotated based on the expression levels of canonical cell-type-specific markers (see **Methods**). In total, we identified 26 clusters across the three groups, each exhibiting differentially expressed genes (DEGs) (**Figure 2I**). These clusters were annotated as ten pancreatic beta-cell subsets, three alpha-cell subsets, four ductal cell subsets, two acinar cell subsets, and three T-cell subsets (**Figure 2J**). Additionally, we identified single clusters corresponding to plasma cells, erythrocyte/immune cells, dendritic cells, and a B-cell population (**Figure 2J**). All ten beta-cell clusters were classified as independent cell types based on their distinct DEGs (**Figure 2J**). Detailed information on cell type descriptions and marker genes for each tissue type is provided in **Tables S1 and S2**.

Using the CellKb^pro^ ranking-based score program (see **Methods**), we annotated all 26 clusters, identifying clusters 10, 14, 20, 21, and 23 as “senescent metaplastic cells” as well as “beta cells” (**Figure 1J**, **Table S2**). Four of the five putative senescent beta-cell clusters (10, 14, 20, and 23) had a higher cell number and proportion (%) in the 30W-S group compared to the 12W-C or 30W-C groups (**Figures 2K**, **S1E**). Gene Ontology (GO) and pathway analyses revealed that clusters 10, 20, 21, and 23 were predominantly enriched in GO terms related to the mitochondrial inner membrane, while cluster 14 was enriched in transport vesicle (**Figure 2L**).

To further confirm whether clusters 10, 14, 20, 21, and 23 represent senescent beta cells, we analyzed each cluster for senescent beta-cell characteristics by comparing the expression of three beta-cell indices: beta-cell identity-related genes, beta-cell disallowed genes, and beta-cell senescence (and aging)-related genes, as previously reported^19,39,70–72^. Research has established that senescent beta cells lose expression of beta-cell identity-related genes while exhibiting increased expression of beta-cell disallowed genes and senescence- and aging-associated genes^19,39,70–72^. We observed that the different senescent beta-cell subsets displayed heterogeneous gene expression levels across the three indices (**Figures 3A**, **S2A**). To more broadly categorize each cluster into different senescence stages, we developed a “*Progression of Senescence (P.S.)*” equation to numerically capture and integrate the diverse genetic markers associated with senescence (see **Methods**). Consistent with the dendrogram analysis (**Figure 3A**), the cell clusters were grouped into three stages using the *P.S.* equation—early, mid, or late. These results suggest that the heterogeneity of senescent beta-cell clusters may reflect “snapshots” of different stages of the senescence process.

**Figure 3.**
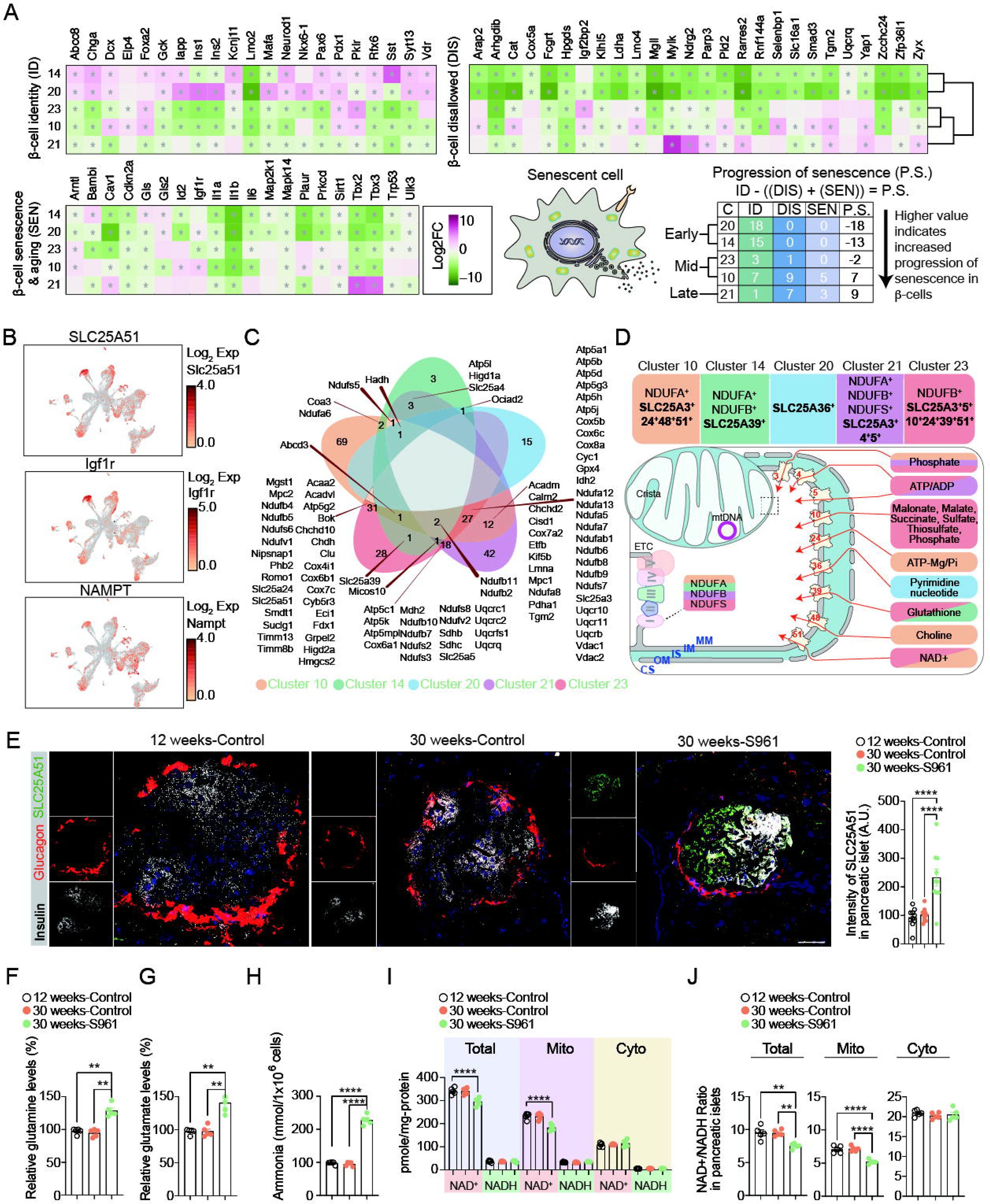
SLC25A51 is part of a senescence gene signature in pancreatic beta cells. Cluster annotation was performed using the CellKB^PRO^ program, as an unbiased approach, to filter out non-beta-cell and non-senescent clusters (**Table S2**). (**A**) The filtered clusters (14, 20, 23, 10, and 21) were further evaluated for senescence characteristics using three different indices: beta-cell identity, beta-cell disallowed, and beta-cell senescence and aging-related genes. Additionally, a dendrogram was constructed to compare each cluster by analyzing the whole scRNA-seq data. Genes with positive Log2FC and p-value (p<0.05) were counted for each index in a cluster to calculate the P.S. Using this method, early-stage senescence (cluster 20, 14), mid-stage (cluster 23, 10), and late-stage (cluster 21) were identified. (**B**) Slc25a51, Igf1r, and Nampt levels were analyzed in UMAP graphs. (**C**) Venn diagram analysis was performed between senescent beta-cell clusters (10, 14, 20, 21, and 23) to identify commonly shared genes. (**D, top box**) Based on the GO:CC of clusters 10, 14, 20, 21, and 23 (mitochondrial inner membrane, transport), these clusters were re-annotated with mitochondrial inner membrane-related genes, including SLC25A family and ETC complex genes. (**D, bottom diagram**) A diagram depicting five different senescent beta-cell clusters, each annotated with its unique differentially expressed mitochondrial inner membrane transporter genes (DEGs). Each DEG is also illustrated in the diagram, showing how different mitochondrial inner membrane transporters facilitate the transport of specific metabolites in different senescent states. Mouse pancreas tissues were stained for insulin, glucagon, and Slc25a51 in (**E**) 12W-C, 30W-C, and 30W-S groups (n=10/group). Scale bars, 50 μm. One-way ANOVA; error bars represent SEM. ****p < 0.0001. (**F-J**) Fresh pancreatic beta cells were isolated from 12W-C, 30W-C, and 30W-S (n=5/group) to analyze (**F**) glutamine, (**G**) glutamate, (**H**) cellular ammonia, (**I**) NAD^+^ and NADH levels, and (**J**) NAD^+^/NADH ratio. One-way ANOVA for comparison between 12W-C, 30W-C, and 30W-S. **p < 0.01, ****p < 0.0001, **p < 0.01, ****p < 0.0001.

To identify commonly shared genes among the different stages of beta-cell senescence, we performed a Venn diagram analysis across five senescent beta-cell clusters. No single DEG was identified as a shared gene across all clusters (**Figure 3C**). However, further analysis identified 89 genes shared between a few clusters, primarily associated with five mitochondrial components: Complex I, Complex III, Complex IV, Complex V, and *SLC25A* family genes (**Figures 3C, 3D, S2B-S2E**). Using these mitochondrial component genes, particularly mitochondrial inner membrane transporter *SLC25A* family genes, we were able to distinguish each cluster: Cluster 10 (*SLC25A3^+^24^+^48^+^51^+^*), Cluster 14 (*SLC25A39^+^*), Cluster 20 (*SLC25A36^+^*), Cluster 21 (*SLC25A3^+^4^+^5^+^*), and Cluster 23 (*SLC25A3^+^5^+^10^+^24^+^39^+^51^+^*) (**Figures 3D, S2C, S2G**). Notably, among the *SLC25A* genes, the mitochondrial NAD^+^-transporter *SLC25A51* was highly expressed in clusters 10, 14, 20, and 23 (**Figures 3B, S2B, S3A-S3C**) and served as a DEG in the ‘mid-stage’ of beta-cell senescence, represented by clusters 10 and 23 (**Figure 3A, 3D**). Overall, SLC25A51 was the sole gene identified as a DEG in more than two senescent beta-cell clusters, and its expression correlated with the beta-cell aging and senescence markers IGF1R and NAMPT, respectively^39,40^ (**Figures 3B, 3C, S2B**).

### SLC25A51 is part of a senescence gene signature in pancreatic beta cells

To confirm whether SLC25A51 protein levels were concomitantly increased in beta cells in the S961 model, we performed IF staining in pancreatic tissues from experimental cohorts. SLC25A51 accumulated in beta cells isolated from S961-treated animals compared to control groups (**Figure 3E**), consistent with our observation in the chronological aging model (**Figure 1E**). To gain a metabolic overview in these mice, we collected serum for targeted metabolomic analysis from both chronologically aged and S961-treated mice. Interestingly, metabolites in pathways influencing NAD^+^ levels or redox balance, including the malate-aspartate shuttle and nicotinate and nicotinamide metabolism, were significantly altered between groups (**Figures S4A-S4F)**. To further understand how these metabolites are altered at the sub-cellular level, we isolated pancreatic islets either from chronologically aged (**Figures 1N, 1O**) and S961-treated mouse models (**Figures 3I, 3J**). In chronologically aged mouse model, we observed a decreased NAD^+^/NADH ratio in whole-cell lysates, cytosol, and mitochondria isolated from pancreatic islets (**Figures 1N, 1O**). However, mitochondrial, cytosolic, and whole-cell NAD^+^ and NADH levels were lower in the S961-treated group, with the NAD^+^/NADH ratio also reduced in the mitochondria and whole-cell lysates, but not in the cytosol, compared to the 12-week-old and 30-week-old untreated control groups (**Figures 3I, 3J**). Similar to the metabolic changes observed in chronological aging (**Figures 1K-1M**), S961-treated mice showed increased glutamine, glutamate, and cellular ammonia levels compared to control groups, indicative of potential changes in glutaminolysis (**Figures 3F-3H**). Glutaminolysis is a metabolic pathway essential for cell survival during senescence, and SLC25A51 is required for glutaminolysis and the entry of glutamate into the TCA cycle^63^. Thus, upregulation of SLC25A51 in pancreatic islets of S961-treated or chronologically aged mice might be an adaptation or a driver of the metabolic changes in beta cells undergoing senescence.

### SLC25A51 is upregulated in senescence downstream of p16^INK4A^ and potentially NRF2, and modulates intracellular NAD^+^ compartmentalization to contribute to the senescence phenotype

To mechanistically probe the functional connection between SLC25A51 and senescence in beta cells, we conducted *in vitro* studies using both murine pancreatic islets and MIN6 mouse insulinoma cells^73^. While SV40 T antigen-immortalized MIN6 cells do not undergo a senescence-associated cell cycle arrest, they recapitulate aspects of senescence, including increased beta-Gal activity and the induction of senescence-associated factors (p16^INK4A^, p21^Cip1/Waf1^, NAMPT) and SASP factors in response to prolonged passaging or H_2_O_2_ treatment ^19,40,74,75^.

Primary pancreatic islet cells isolated from 30-week-old C57BL/6N mice (n=5/group) exhibited increased beta-Gal activity in a concentration-dependent manner following H_2_O_2_ treatment, peaking at 200 μM, with diminished activity and increased cell death at concentrations exceeding 500 μM (**Figures 4A, S5A,** data not shown). Thus, we evaluated mRNA and protein levels of senescence markers 24 hours after treatment with 200 μM H_2_O_2_. H_2_O_2_ treatment elevated SASP production (CXCL10, IL-6, and IL-1β) and increased mRNA levels of *Slc25a51* and the beta-cell senescence markers *Igf1r*, *Cdkn2a*, *Trp53*, and *Il-1b* (**Figures 4A-4C, S5A**). Additionally, IGF1R and SLC25A51 protein levels were increased in pancreatic islet cells following H_2_O_2_ treatment (**Figure 4D**). Similar to pancreatic islet cells, SLC25A51 and senescence markers (p16^INK4A^, p21^Cip1/Waf1^, γ-H2AX, NAMPT) were upregulated in a concentration- and time-dependent manner in MIN6 cells, with maximal levels observed at 200 µM for 24 hours (**Figures 4F**). Treatment with MitoTempo, a mitochondria-localized redox modulator that removes reactive oxygen species (ROS) and reduces diabetic complications^76^, prevented the H_2_O_2_-induced upregulation of SLC25A51 and senescence markers, including p16^INK4A^, p21^Cip1/Waf1^, NAMPT, and γ-H2AX (**Figure 4K**). Oxidative stress induced by H_2_O_2_ treatment not only triggers senescence but also impairs glucose-induced insulin secretion in pancreatic beta cells by depolarizing the mitochondrial membrane potential^77^. Indeed, TMRE and mitoSOX staining revealed that H_2_O_2_ decreased mitochondrial membrane potential and increased mitochondrial superoxide, respectively (**Figure 4E**), without altering mitochondrial mass as indicated by MitoTracker staining (**Figure 4E**). While citrate synthase and Complex I proteins increased following H_2_O_2_ treatment, MTCO2 and TOM20 levels remained unchanged (**Figures 4F, S5B**). Because H_2_O_2_ treatment directly impacts mitochondrial physiology, we further investigated whether the mitochondrial NAD^+^ transporter SLC25A51 was influenced by replicative senescence or irradiation with 10 Gy of ionizing radiation (IR). Similar to H_2_O_2_ treatment, both prolonged passaging and irradiation of MIN6 cells increased SLC25A51 levels and senescence markers such as p16^INK4A^, p21^Cip1/Waf1^, and γ-H2AX (**Figures 4I, 4J**). These results indicate that SLC25A51 accumulation occurs in response to beta-cell senescence in a beta-cell-specific, oxidative stress/ROS-dependent manner.

**Figure 4.**
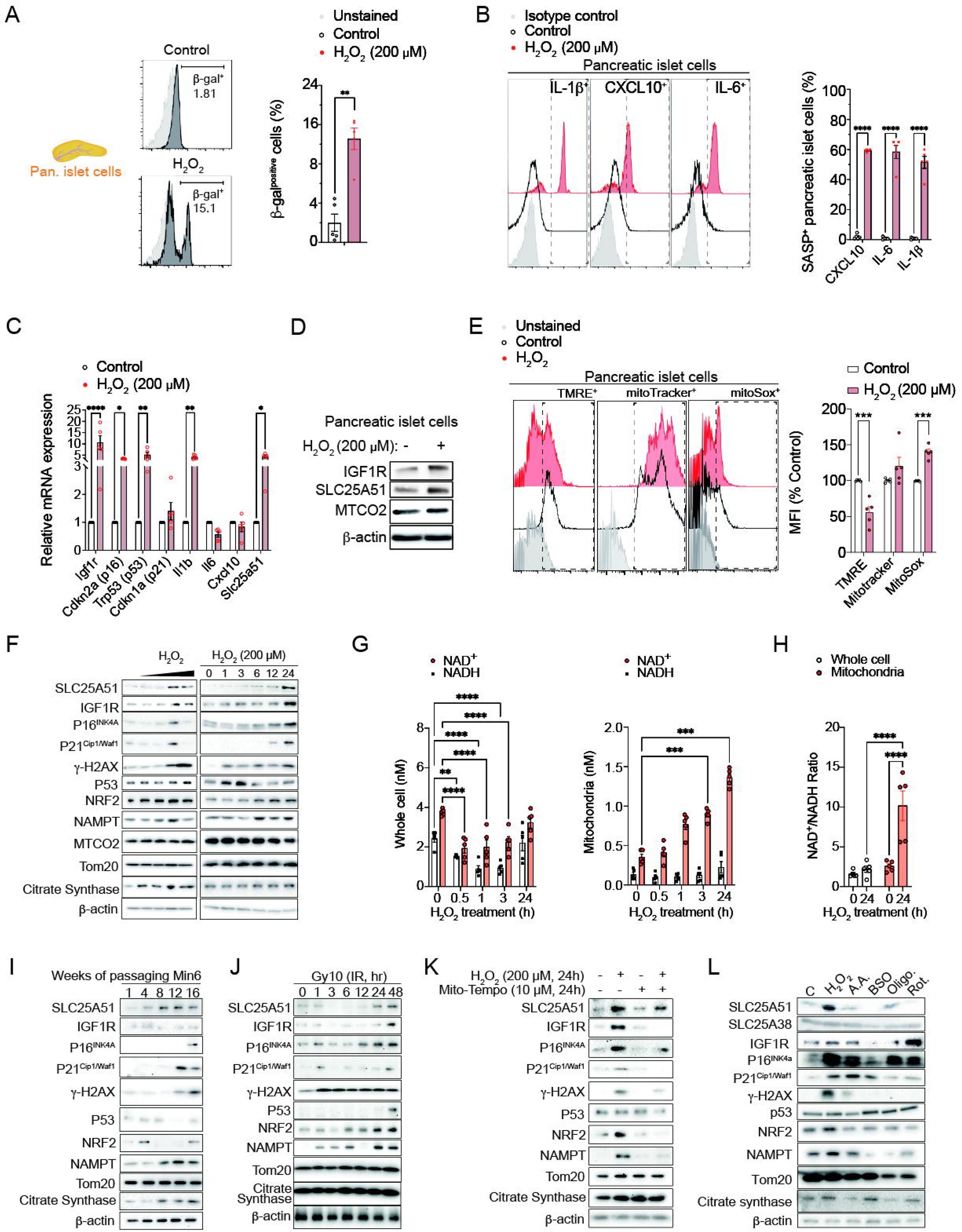
SLC25A51 is upregulated in senescence downstream of the senescence-associated cell cycle regulator p16INK4A, and modulation of intracellular NAD compartmentalization by SLC25A51 contributes to the senescence phenotype. (**A-E**) Thirty-week-old C57BL/6N male mice were sacrificed to isolate pancreatic beta cells and cultured with either PBS (control) or H_2_O_2_ (200 μM, 24 hours) (n=5/group). Then, (**A**) beta-Gal activity, (**B**) SASP, (**C**) mRNA levels of senescence markers (*Igf1r, Cdkn2a, Trp53, Cdkn1a, Il1b, Il6, Cxcl10*) and *Slc25a51* were analyzed (n=5/group). (**D**) Protein levels of IGF1R, SLC25A51, MTCO2, and β-actin were analyzed. Further€e, (**E**) mitochondrial membrane potential (TMRE), MitoTracker, and MitoSOX were analyzed (n=5/group). Statistical analysis was performed using either (**A**) two-tailed *t*-test or (**B, C, E**) two-way ANOVA; error bars represent SEM; *p < 0.05, **p < 0.01, ***p < 0.001, ****p < 0.0001 for comparison between control (PBS) and H_2_O_2_-treated groups. (F) SLC25A51 expression levels and other protein levels were analyzed in MIN6 cells with replicative senescence by treating with H_2_O_2_ in a concentration-dependent manner with 0, 50, 100, 200, and 500 μM, or in a time-dependent manner (hour). At least three repeats of Western blot were performed for each experiment. (**G, H**) MIN6 cells overexpressed with pMXs-HA-Mito-GFP were generated to analyze mitochondrial NAD^+^ and NADH levels. Cells were treated with either control (PBS) or H_2_O_2_ (200 μM) in a time-dependent (0, 0.5, 1, 3, 24 hours) manner. Then, fresh whole-cell lysates and isolated mitochondria were obtained from each group to analyze (**G**) NAD^+^ and NADH levels at 0, 0.5, 1, 3, and 24 hours (n=5/group) and (**H**) NAD^+^/NADH ratio at 0 and 24 hours, measured by a nanomolar-sensitive NAD^+^/NADH-Glo assay. Statistical analysis was performed using two-way ANOVA; error bars are SEM, *p < 0.05, **p < 0.01, ***p < 0.001, ****p < 0.0001. (I) SLC25A51 and other protein levels were analyzed in MIN6 cells passaged for (**G**) 1, 4, 8, 12, and 16 weeks. At least three trials of Western blot were performed. (J) Six cells were irradiated with 10 Gy of ionizing radiation (IR) and analyzed at the indicated time points for SLC25A51 and other protein levels. At least three trials of Western blot were performed. (K) Six cells were treated with or without H_2_O_2_ (200 μM) in the presence or absence of Mito-Tempo (10 μM) to assess SLC25A51 and other protein levels. At least three trials of Western blot were performed. (L) Six cells were treated with either H_2_O_2_ (200 μM), Antimycin A (A. A., 0.5 μM, 24 hours), BSO (1 mM, 24 hours), Oligomycin A (Oligo., 0.5 μM, 24 hours), or Rotenone (Rot., 0.5 μM, 24 hours) to assess SLC25A51 and other protein levels. At least three trials of Western blot were performed.

SLC25A51 is the major mitochondrial NAD^+^ transporter in mammals^50^, and it is conceivable that its upregulation is a consequence of the drop in whole-cell or mitochondrial NAD^+^ levels or the NAD^+^/NADH redox ratio in pancreatic islets that we and others have observed during aging and senescence^3-5,7,31,78,79^ (**Figures 1O, 1P, 3I, 3J**). In MIN6 cells cultured in media containing excess concentrations of NAD^+^ precursors, such as nicotinamide, we observed a decline in total cellular NAD^+^ levels during H_2_O_2_-induced senescence from 0.5 to 3 hours, followed by a re-stabilization of NAD^+^ levels at 24 hours (**Figure 4G**). The decrease in total cellular NAD^+^ levels and increased oxidative stress from H_2_O_2_ at early time points led to the induction of NRF2, a master regulator of the oxidative stress response, between 6 and 24 hours (**Figure 4F**). NRF2 activation can upregulate the NAD^+^-consuming enzyme ALDH3A1 and decrease the NAD^+^/NADH ratio in lung cancer cells^80^. Whether NRF2 directly regulate *Slc25a51* has not been evaluated. Analysis on the upstream regions of the human and mouse *Slc25a51* sequences revealed potential antioxidant response element (ARE) and p53-response element (p53-RE) sequences (**Figures S4G**). In our scRNA-seq analysis, we observed that the *Nfe2l2* (NRF2), *Nfe2l1* (NRF1), and *Trp53* (p53) genes were upregulated in clusters with high *Slc25a51* expression levels (**Figures S2B, S4K**). Furthermore, transcription factor prediction programs (JASPAR^81^ and PROMO^82^) identified *Nfe2l1* (NRF1) and *Nfe2l2* (NRF2) as candidates for regulating *Slc25a51* expression (**Figures S4G-S4I**). By analyzing published data from a genome-wide CRISPR screen in KEAP1-NRF2 interaction inhibitor (KI696)-treated KEAP-1-dependent CALU6 cells^80^, we found that *Slc25a51* was one of the genes sensitive to NRF2 activation (**Figure S4J**). Furthermore, H_2_O_2_-treatment, irradiation, and overexpression of *Nfe2l2* in MIN6 cells led to an increase in SLC25A51 levels (**Figures 4F, 4J, 5A**). These data implicate NRF2 as a potential regulator of SLC25A51 expression in beta cells, suggesting a mechanism to induce mitochondrial NAD^+^ influx and metabolic adaptation in response to oxidative stress. To assess whether H_2_O_2_ treatment indeed induces mitochondrial NAD^+^ influx, we isolated mitochondria to measure NAD^+^ and NADH levels. Unlike total NAD^+^ levels, H_2_O_2_ treatment increased mitochondrial NAD^+^ levels ∼4-fold after 24 hours and the NAD^+^/NADH redox ratio ∼3-fold, concurrent with an increase in SLC25A51 protein levels, while whole-cell NAD^+^ and NADH levels remained unchanged (**Figures 4G, 4H**, **S5G-S5H**). In contrast, treatment with the Complex I inhibitor, Rotenone (Rot), or other electron transport chain (ETC) inhibitors (Antimycin A [A.A] and Oligomycin A [Oligo]) did not induce SLC25A51 or NRF2 expression (**Figures 4L, S5B**). Induction of oxidative stress by treating cells with the glutathione synthesis inhibitor BSO increased senescence markers, such as p16^INK4A^ and p21^Cip1/Waf1^, but failed to induce SLC25A51 or NRF2 (**Figure 4L**). Moreover, reducing whole-cell and mitochondrial NAD^+^ by inhibiting the NAD^+^ salvage enzyme NAMPT with FK866 did not induce SLC25A51 expression (**Figures S5F-S5H**).

Thus, we concluded that SLC25A51 upregulation does not occur as a compensatory response to lower cellular or mitochondrial NAD^+^ levels, changes in the NAD^+^/NADH redox ratio, or mitochondrial dysfunction. Instead, it may serve as part of the senescence response to drive metabolic changes, such as increased glutaminolysis, that are necessary to sustain the senescent cell state.

### SLC25A51 overexpression induces senescence markers in MIN6 cells

While senescence can be induced downstream of DNA damage, metabolic or mitochondrial changes such as impaired glucose sensing and insulin production in beta cells^18^, and mitochondrial insults^24,28,29,83,84^ in different senescent cell types, are central steps in the senescence program. To determine whether SLC25A51 induction occurred as part of the senescence program, we tested whether overexpression of senescence factors or the putative transcriptional regulator NRF2, in the absence of other senescence stimuli, was sufficient to induce SLC25A51. Overexpression of p16^INK4A^, and to a lesser degree p21^Cip1/Waf1^ and NRF2, was indeed sufficient to induce SLC25A51 protein (and other senescence markers) in MIN6 cells (**Figure 5A**), while levels of another mitochondrial transporter, SLC25A38, remained unchanged. In addition, overexpression of SLC25A51, which led to a more than 3-fold increase in mitochondrial NAD^+^ levels, and a 2-fold increase in the mitochondrial NAD^+^/NADH ratio, but not its transport-dead K91A point mutant^50,85^ or SLC25A38^86^, was sufficient to induce p16^INK4A^, p21^Cip1/Waf1^, and γ-H2AX (**Figures 5E-5G**). Thus, we reasoned that SLC25A51 upregulation downstream of p16^INK4A^, possibly through its role in compartmentalizing NAD^+^ and controlling its redox balance, modulates senescence or could be part of a feed-forward loop that further drives the senescence program.

**Figure 5.**
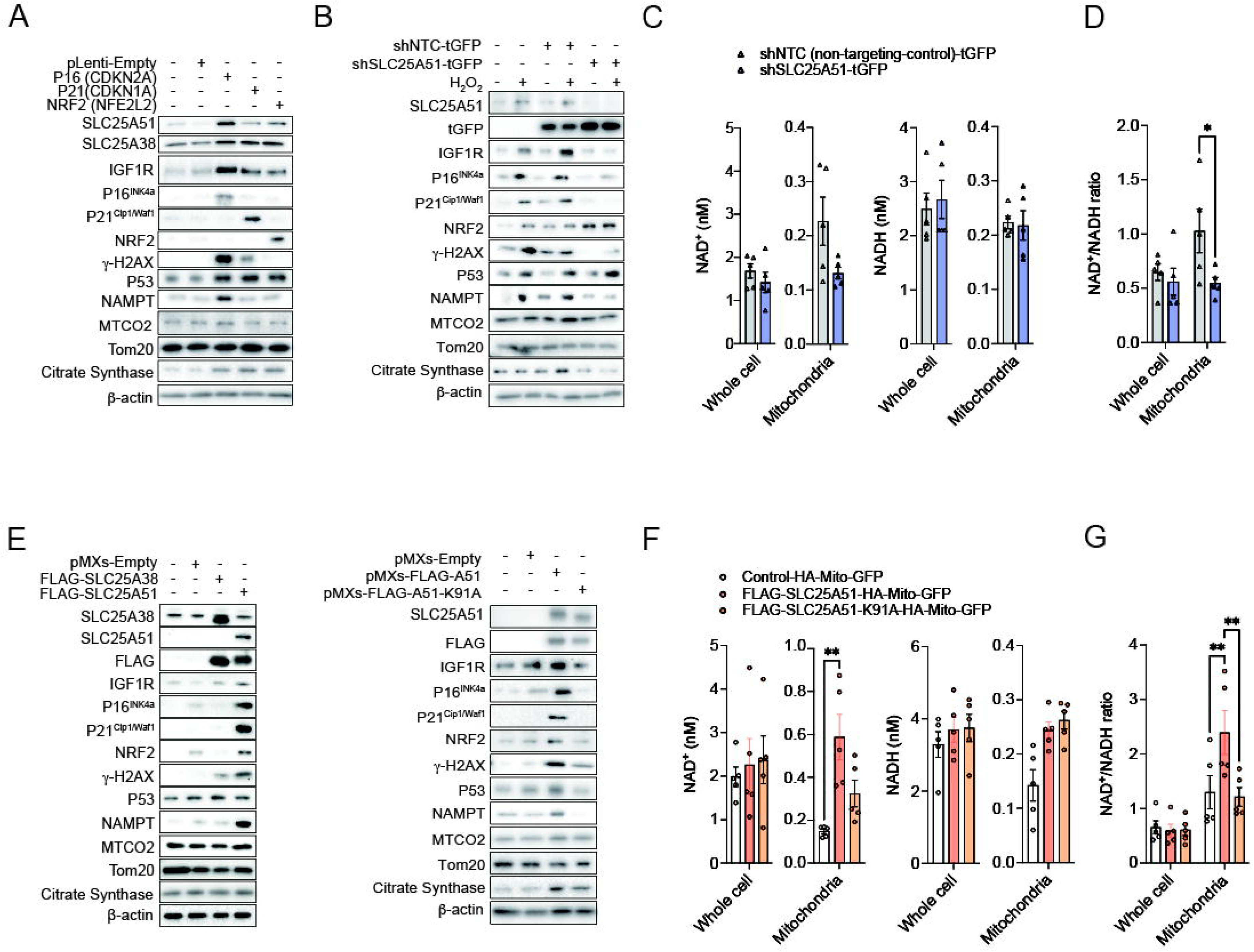
SLC25A51 overexpression induces senescence markers in MIN6 cells. MIN6 cells were stably transfected with (**A**) pLenti-Empty, pLenti-*Cdkn1a* (p21^Cip1/Waf1^), pLenti-*Cdkn2a* (p16^INK4A^), or pLenti-*Nfe2l2* (Nrf2), (**B**) pLKO.1-shEmpty-tGFP or pLKO.1-shSLC25A51-tGFP treated with or without H_2_O_2_ (200 μM, 24 hou€, and (**E**) pMXs-Empty, pMXs-Flag-*Slc25a51*, or pMXs-Flag-*Slc25a38*. (**A, B, E**) Whole-cell lysates from each condition were probed for SLC25A51, IGF1R, p16^INK4A^, p21^Cip1/Waf1^, p53, γ-H2AX, NRF2, MTCO2, TOMM20, citrate synthase, and β-actin. Additionally, (**A, E**) Flag, SLC25A38, or (**B**) tGFP were tested. At least three trials of Western blot were performed. (**C, D, F, G**) Fresh whole-cell lysates and mitochondria were isolated from each group to analyze NAD^+^, NADH, and the NAD^+^/NADH ratio in (**F, G**) HA-mito-GFP overexpressed pMXs-Empty, Flag-Slc25a51, or Flag-Slc25a51-K91A-mutant cells and (**C, D**) pLKO.1-shEmpty-tGFP or pLKO.1-shSLC25A51-tGFP cells. Statistical analysis was performed using two-way ANOVA; error bars are SEM, *p < 0.05, **p < 0.01.

If metabolic changes downstream of SLC25A51’s role in compartmentalizing NAD^+^ or controlling the cellular NAD^+^/NADH ratio contribute to the senescence program, we hypothesized that SLC25A51 depletion, in turn, could prevent senescence induction. Indeed, SLC25A51 depletion in MIN6 cells using stable shRNA transfection, which led to a reduction in mitochondrial NAD^+^, prevented H_2_O_2_-induced upregulation of senescence markers (p16^INK4A^, p21^Cip1/Waf1^, and γ-H2AX) (**Figures 5B-5D**). This finding aligns with a recent study in mammalian U2OS cells, where SLC25A51 knockdown prevented H_2_O_2_-induced DNA damage by increasing nuclear availability of NAD^+^ for PARP1-dependent DNA repair and phosphorylation of histone γ-H2AX, a marker of DNA damage and senescence^87,88^.

### Beta cell-specific deletion of SLC25A51 improves age-associated glucose homeostasis and insulin resistance in mice

Chronological aging and metabolic insults induce pancreatic beta-cell senescence and insulin resistance in mice and humans^6,18,19,23,39,60,77,84,89–102^. The accumulation of senescent beta cells and the increased risk of insulin resistance during aging contributes to T2D pathogenesis through both cell-autonomous and non-autonomous mechanisms^19,27,38–41,101,103^. Developing strategies to prevent beta-cell senescence could have major implications for the treatment of T2D^19,39^. Given that SLC25A51 is a key regulator of mitochondrial NAD^+^ metabolism and its levels correlate with increased senescence and inflammation, we further investigated whether inhibiting SLC25A51 in insulin-resistant male mice could prevent age-associated diabetes.

As in mice, SLC25A51 was predominantly expressed in beta cells in the human pancreas (**Figure S1B**). To assess SLC25A51’s potential as a therapeutic target, we analyzed SLC25A51 expression in scRNA-seq datasets from humans. *SLC25A51* levels were higher in male T2D patients compared to healthy male controls from two different datasets, similar to the expression of *Cdkn1a* (p16^INK4A^) and *Cdkn2a* (p21^Cip1/Waf1^) (**Figures 6A, S6B, S6C**)^104,105^. To test the effect of SLC25A51 deletion on glucose homeostasis and insulin sensitivity, we specifically deleted SLC25A51 in beta cells, employing a SLC25A51*floxed* mouse model we developed (**Figures 6A, S6A, S6E**). Using AAV6 viruses harboring Cre recombinase driven by the beta-cell-specific insulin 1 promoter (INS1-Cre), we selectively deleted SLC25A51 in pancreatic beta cells of 30-week-old male and female mice. Control mice received AAV6 viruses with an empty vector (INS1-Empty) or PBS via pancreatic duct injection (**Figures 6A, S6A, S6E**). Fifteen weeks post-surgery, male mice were sacrificed for pancreatic islet and tissue analysis. Western blot and IF staining confirmed the loss of SLC25A51 protein in the pancreatic islets of INS1-Cre-infused mice compared to INS1-Empty mice (**Figures 6I, S6D**). Furthermore, beta-cell-specific SLC25A51 knockout (βKO) mice exhibited reduced p16^INK4A^ levels, indicating lower beta-cell senescence (**Figures 6J, S6D**). Since age-related insulin resistance is observed in both mice and humans, we further investigated whether the loss of SLC25A51 in pancreatic beta cells affected insulin sensitivity. While infusion of INS1-CRE AAV6 viruses resulted in a slight reduction in body mass and blood glucose levels compared to the INS1-Empty AAV6 and PBS-infused mice (**Figures 6B, 6D**), circulating beta-hydroxybutyrate levels remained unchanged (**Figure 6C**). Interestingly, blood glucose levels in SLC25A51 βKO mice remained lower after virus infusion, while body weight stabilized to pre-surgery levels (**Figure 6D**).

**Figure 6.**
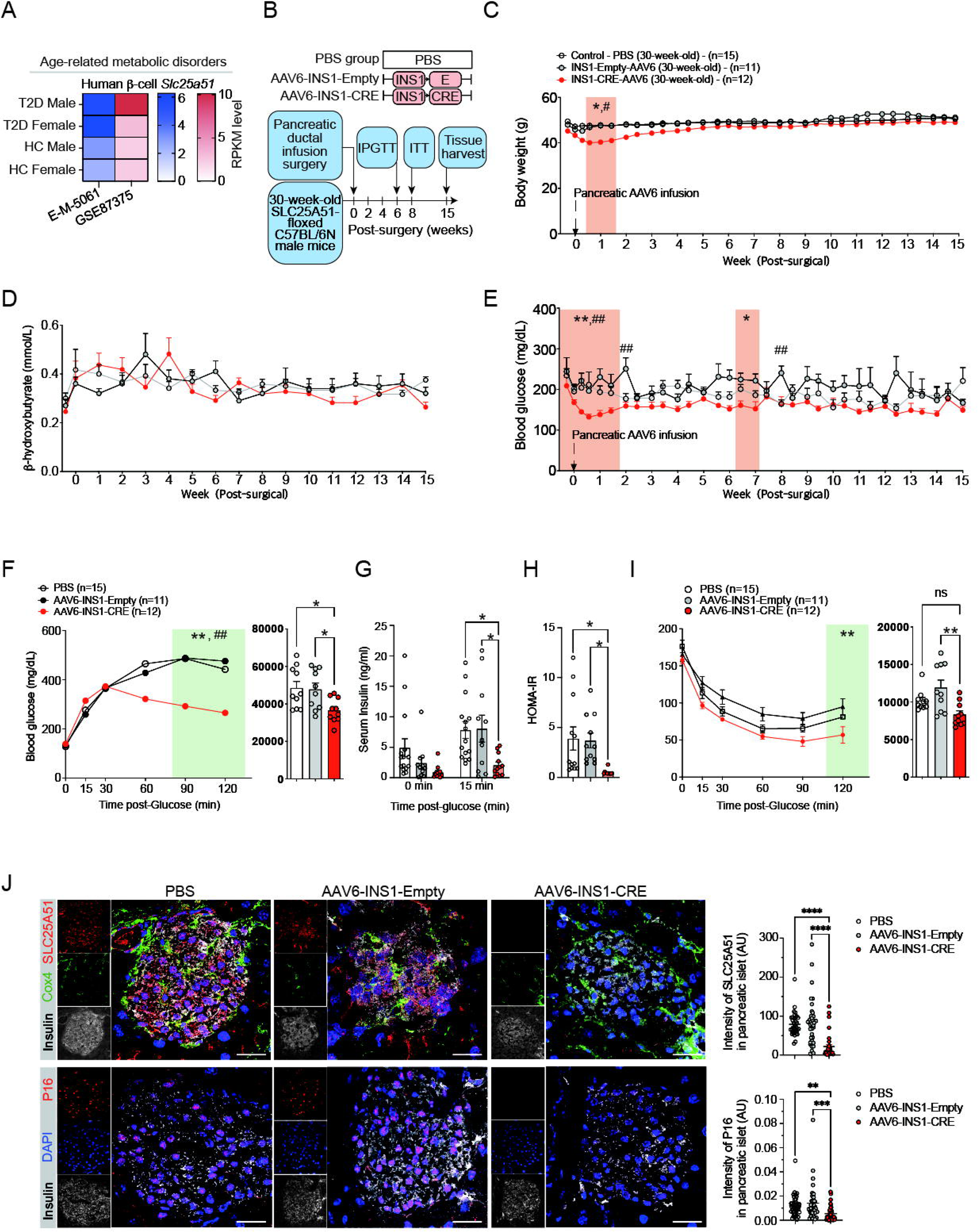
Beta cell-specific deletion of SLC25A51 improves age-associated glucose homeostasis and insulin resistance in mice. (A) Human *Slc25a51* levels were analyzed in T2D patients and healthy controls from publicly available datasets (E-M-5061, GSE87375). (B) The diagram depicts the strategy for infusing AAV6-INS1-CRE viruses through the pancreatic duct to specifically knock out beta-cell-specific SLC25A51 in 30-week-old SLC25A51 floxed C57BL/6N mice. After infusion and recovery, mice were analyzed for body weight, random blood glucose levels, β-hydroxybutyrate levels, IPGTT, and ITT under specified conditions and sacrificed after 15 weeks. (**C-E**) Pancreatic ductal infusion of either PBS, AAV6-INS1-Empty, AAV6-INS1-CRE was performed in each group. At 15 weeks after surgery, pancreatic islet cells and pancreatic tissues were isolated to confirm the knockout of SLC25A51 (**Figure S6D**) using Western blots and (**J**) I€taining. (**C**) Body weight and (**D**) random blood glucose levels were analyzed twi€a week, and (**E**) β-hydroxybutyrate levels were analyzed once a week for each mouse. Statistical analysis was performed using repeated measurement two-way ANOVA; error bars represent SEM; *p < 0.05, **p < 0.01 between PBS and AAV6-INS1-CRE groups, ^#^p < 0.05, ^##^p < 0.01 between AAV6-INS1-Empty and AAV6-INS1-CRE groups. (**F-I**) Six or eight weeks after surgery, mice were subjected to IPGTT and ITT. Graphs show blood glucose levels at (**G**) each time point and area under the curve. During IPGTT, serum was collected from each group at 0 and 15 min to measure (**H**) serum insulin. After two weeks from IPGTT, (**I**) ITT was performed to test insulin sensitivity. Two-way ANOVA; error bars represent SEM; **p < 0.01 comparison between PBS and AAV6-INS1-CRE, ^##^p < 0.01 comparison between AAV6-INS1-Empty and AAV6-INS1-CRE. See also **Figures S6A-S6G** for female data. HOMA-IR was calculated for each group in male mice. HOMA-IR values between 0.5 and 1.4 are considered normal, ≥ 1.9 is regarded as early insulin resistance, and above 2.9 indicates insulin resistance. (J) After sacrifice, mouse pancreatic islets were acquired from each group. IF staining of (up) DAPI, insulin, Cox4, and Slc25a51, or (down) DAPI, insulin, and p16^INK4A^ were performed in pancreatic sections. Pancreatic sections were prepared as described; representative data are from 2–5 pancreatic islets per mouse in each group (n=35∼40 islets/group). Scale bars, 20 μm. Two-way ANOVA; error bars are SEM. **p<0.01, ***p<0.001, ****p<0.0001.

To assess glucose clearance and insulin sensitivity, we performed the intraperitoneal glucose tolerance test (IPGTT) and insulin tolerance test (ITT). SLC25A51 βKO mice displayed improved glucose clearance in the IPGTT. Insulin measurements at 0 and 15 min during IPGTT and ITT revealed hyperinsulinemia in the two control groups, while SLC25A51 βKO mice exhibited lower systemic insulin levels (PBS:7.84±5.2 ng/ml; AAV6-INS1-empty:8.07±7.3 ng/ml; AAV6-INS1-CRE:2.07±1.8 ng/ml), likely explaining the improved insulin sensitivity (**Figures 6E-6H**). HOMA-IR analysis indicated insulin resistance in the PBS and AAV6-INS1-Empty groups, whereas the AAV6-INS1-CRE infused group was non-insulin-resistant (**Figure 6G**). Like male mice, female mice also showed improved insulin sensitivity (**Figure S6J**). However, glucose levels, serum insulin levels, and IPGTT showed minimal effects in female mice (**Figures S6F-S6I, S6K**). Furthermore, IF staining suggested that the improved metabolic phenotype in male mice upon beta-cell-specific SLC25A51 deletion was due to lowered levels of p16^INK4A^ -positive senescent beta cells (**Figures 6I, J**).

## Discussion

Pancreatic beta cells are the primary insulin-producing cells, essential for controlling systemic glucose homeostasis. As postmitotic cells, pancreatic beta cells are irreplaceable or rarely replaced with age, and the accumulation of senescent beta cells during aging is detrimental to organismal health^19,27,38–41,101,103^. This accumulation partially contributes to the loss of beta cell function and/or mass, a hallmark of both T1D and T2D^19,27,39–41,101^. Heterogeneity in islet and senescent beta cell profiles, along with a lack of experimentally tractable models that faithfully recapitulate beta-cell biology, have complicated the study of the mechanisms underlying beta-cell senescence and dysfunction. Our work highlights the central role of NAD^+^ compartmentalization and redox balance in driving cell state and fate.

SLC25A51 is expressed in various tissues and cell types, and we are beginning to uncover its diverse functions in metabolic regulation through its role in mitochondrial NAD^+^ transport and intracellular compartmentalization. Increased SLC25A51 levels correlate with elevated tumor growth or cancer cell proliferation. In acute myeloid leukemia (AML) patients, increased SLC25A51 levels are associated with poor survival outcomes, and lowering SLC25A51 levels prevented tumor growth in U937-engrafted female mice^45^. Similar observations have been made in other types of cancer^87,106^. In human thoracic aortic aneurysm, SLC25A51 expression levels inversely correlate with disease severity and postoperative progression^107^. Low SLC25A51 expression increases the risk of aortic aneurysm and dissection^107^. In mouse models, SLC25A51 deletion caused the most severe aortic aneurysm compared to other smooth muscle-specific knockouts of Nampt, Nmnat1, Nmnat3, Nadk2, and Aldh18a1^107^. However, the role of SLC25A51 in aging and senescence has not been studied.

As a mitochondrial NAD^+^ transporter, SLC25A51 plays a key role in regulating intracellular NAD^+^ distribution and redox ratio. Its activity influences not only mitochondrial NAD^+^ levels, but also nucleo-cytoplasmic NAD^+^ distribution in certain cell types^87^. This regulation affects processes such as the epigenetic state, DNA repair, and metabolic function through sirtuins, PARPs, and AMPK, as well as cell cycle regulation and senescence through p53, p16^INK4A^, and p21^Cip1/Waf1108^. Modulating mitochondrial NAD^+^ transporters such as Ndt1/Ndt2 has been shown to extend chronological lifespan in yeast^48^ and Arabidopsis plants^109^ by enhancing mitochondrial respiration efficiency and reducing ROS production. Notably, recent studies on beta-cell aging found that NAD^+^-related genes (Nrk1, Nmnat2) are upregulated with aging and disrupt the intracellular NAD^+^/NADH ratio^4,43^. Our results imply that mitochondrial NAD^+^ transporter induction is an evolutionarily conserved feature of chronological aging in tissues or cell types exposed to oxidative stress and metabolic insults. Furthermore, the accumulation of SLC25A51 in senescent beta cells is not merely a cellular adaptation to decreased mitochondrial NAD^+^ levels but may actively contribute to modifying senescence. This could occur by modulating mitochondrial or nuclear NAD^+^ levels and affecting downstream NAD^+^-dependent enzymes such as Sirt1 and PARP1 or uncoupling the NAD^+^/NADH ratio. Hyperglycemia is an important driver of senescence, cell identity loss, and beta-cell dysfunction, though the mechanism remains unclear^28^. Increased glycolytic rates have been shown to drive adaptation in NAD^+^ compartmentalization, affecting beta cell identity and function^43^. In rapidly proliferating cancer cells, SLC25A51 is upregulated to decouple mitochondrial NAD^+^/NADH, providing a proliferative advantage by supporting oxidative reactions from a variety of fuels^45^. Like cancer cells, senescent cells shift their metabolism to support survival, including increased ammonia production to counteract an acidic intracellular milieu^63^. It is interesting to speculate that in the context of hyperglycemia, SLC25A51 knockdown may counteract the increased glycolytic flux driving senescence or the metabolic changes required for adaption to the senescent cell state.

The mechanisms through which SLC25A51 depletion counteracts senescence remain unclear. In a breast cancer model, SLC25A51 depletion increased nuclear NAD^+^ levels and thus enhanced PARP1-dependent DNA repair^47^. Whether this mechanism also operates in beta cells and how SLC25A51-mediated regulation of NAD^+^-dependent DNA repair and chromatin remodeling enzymes affects beta-cell function and epigenetic state remain to be elucidated. Of note, mitochondrial coupling of the NAD^+^/NADH ratio is directly linked to ATP production and mitochondrial membrane hyperpolarization in controlling the secretion of insulin granules from beta cells. ATP is subsequently transferred to the cytosol, raising the ATP/ADP ratio and triggering the closure of K_ATP_ channels. This depolarizes the beta-cell membrane, opening voltage-dependent Ca^2+^ channels, increasing cytosolic Ca^2+^, and triggering insulin secretion^110^. Other factors such as ATP, GTP, NADPH, cAMP, LC-CoAs, and glutamate can also promote insulin exocytosis^110^. Future research should explore the acute and chronic effects of SLC25A51 inhibition on beta-cell biology and function. Additional questions include understanding the mechanisms driving SLC25A51 upregulation, identifying the signals and transcriptional programs involved, and investigating the metabolic and signaling pathways regulated by SLC25A51 activity in the context of senescence. Determining whether these features are universal across senescent cells or specific to beta cells, and how SLC25A51 expression relates to other features of the highly heterogenous senescent beta cell populations is also essential. Lastly, studies are needed to clarify how SLC25A51 deletion in beta cells improves systemic insulin sensitivity and glucose homeostasis. Linking mechanistic insights from *in vitro* cell models to physiological observations *in vivo* will yield valuable insights.

## Methods

### Animal Housing

Animal experiments were conducted in the specific pathogen-free environment at the animal facility at Seoul National University Hospital with approval from the Institutional Animal Care and Use Committee (IACUC no. 21-0089-S1A0) (**Figures 1E-1O, 2A-2L, 3A-3J, 4A-4D, S1C-S1G, S2A-S2E, S3A-S3C, S4A-S4F, S4K, S5A**) and at the Harvard Center for Comparative Medicine, Harvard Medical School, Harvard University with approval from the IACUC (no. IS00003204) (**Figures 1B-1D, 4E, 6B-6J, S6A, S6D, S6E-S6K**). Mice were maintained on a 12-hour light/dark cycle and were fed PicoLab Mouse Diet 20 (5058), formulated with 20% protein and 9% fat.

### C57BL/6N Mice

To avoid confounding effects from the loss of nicotinamide nucleotide transhydrogenase (*Nnt),* which could contribute to differences in metabolic phenotype, *Nnt*-wild-type C57BL/6N mice were employed instead of C57BL/6J mice, which have a known mutation in *Nnt* exon 7–11^92^. Twelve-, 30-, and 90-week-old C57BL6/N male mice were acquired from Joongah Bio (Korea) or from wild-type littermates produced from SLC25A51 heterozygote floxed mice. Random blood glucose levels (*ad libitum*) and body weight were measured twice a week for SLC25A51-floxed mice infused with either PBS, AAV6-INS1-Empty, or AAV6-INS1-CRE (**Figure 6**). For S961-treatment experiments, random blood glucose levels and body weight were measured once a day for 14 days (**Figure 2**). Circulating β-hydroxybutyrate levels were measured once a week for SLC25A51-floxed mice infused with PBS, AAV6-INS1-Empty, or AAV6-INS1-CRE (**Figure 6**). After sacrifice, pancreatic tissues were collected from mice, stored in optimal cutting temperature (OCT) compound, or fixed in 4% paraformaldehyde for further analyses. Isolation of pancreatic islet cells, splenocytes, and hepatocytes from C57BL/6N mice was performed as described below.

### Generation of SLC25A51 fl/fl C57BL/6N Mice

*Slc25a51* floxed C57BL/6N mice were generated using a CRISPR/Cas9-based approach by Biocytogen Pharmaceuticals. Briefly, two single-guide RNAs (sgRNAs) were designed using the CRISPR design tool (http://www.sanger.ac.uk/htgt/wge/) to target regions upstream of the exon 3 and downstream of the 3’UTR. These were screened for on-target activity using a Universal CRISPR Activity Assay (UCA^TM^, Biocytogen Pharmaceuticals). To minimize random integrations, a circular donor vector was used. The gene targeting vector, containing a 5’ homologous arm, a target fragment (part of the 5’UTR, exon 3 coding region, and 3’UTR), and a 3’ homologous arm, served as a template to repair the double-strand breaks generated by Cas9/sgRNA. Two LoxP sites were precisely inserted on both sides of the target fragment of the *Slc25a51* gene. T7 promoter sequence was added to the Cas9 or sgRNA template via PCR amplification *in vitro*. Cas9 mRNA, the targeting vector, and sgRNAs were co-injected into the cytoplasm of one-cell-stage fertilized C57BL/6N eggs. The injected zygotes were transferred into the oviducts of Kunming pseudo-pregnant females to generate F0 mice. F0 mice with the expected genotype, confirmed by tail genomic DNA PCR and sequencing, were mated with C57BL/6N mice to establish germline-transmitted F1 heterozygous mice. F1 heterozygous mice were genotyped using tail genomic PCR, Southern blot, and DNA sequencing. Two primer sets were used to confirm SLC25A51 LoxP sites. Primer (F0): AATAAAAGGTGGTTCATAGGGTGGT, (R0): TACTGAAACAGGGGCCACTAAGAT. Primer (F1): AAGAACAGGGATATCGGCCGGGCGT, (R1): AGCCATCTCTTACAGGACCTCTACT.

### AAV6 Virus Production, Infusion, and Surgical Procedures

The rat INS1 promoter sequence, as described previously^111^, was inserted into an AAV6-CRE backbone, which is commonly used for pancreatic ductal infusion^112–114^. Subsequently, AAV6-INS1-Empty (1.81×10^13^ GC/ml) and AAV6-INS1-CRE (7.16×10^13^ GC/ml) were produced by the Penn Vector Core, University of Pennsylvania School of Medicine. For the pancreatic ductal infusion of AAV6 in SLC25A51 fl/fl mice, we infused 10^12^ GC/ml per mouse (30-week-old mouse body weight: 45∼50 g, infusion rate: 6 μl/min, volume: 180 μl), as described by Xiao et al^113^.

### scRNA-seq of Mouse Pancreatic Islet Cells

Chromium Next GEM Single Cell 3’ v3.1 was used for scRNA-seq on mouse pancreatic islet cells (Deposited data: https://osf.io/yaxd2/?view_only=115cd380bf5246a6a77ce3d38049ded2). Libraries were prepared using the Chromium controller according to the 10x Chromium Next GEM Single Cell 3’ v3.1 protocol (CG000204). Briefly, the cell suspensions were diluted in nuclease-free water to achieve a targeted cell count of 5,000. The cell suspension was mixed with a master mix and loaded with Single Cell 3’ v3.1 Gel Beads and Partitioning Oil into a Chromium Next GEM chip G. RNA transcripts from single cells were uniquely barcoded and reverse-transcribed within droplets. cDNA molecules were pooled and underwent end repair, the addition of a single ‘A’ base, and adapter ligation. The products were purified and enriched by PCR to create the final cDNA library. The purified libraries were quantified using qPCR according to the qPCR Quantification Protocol Guide (KAPA) and qualified using the Agilent Technologies 4200 TapeStation (Agilent Technologies). Libraries were then sequenced using the HiSeq platform (Illumina). Read lengths used were 28 for Read1 (Cell barcode & UMI length), 10 for i7 and i5 index (sample index), and 90 for Read2 (insert).

### Evaluating the Progression of Senescence for Beta-Cell Clusters

The filtered clusters (14, 20, 23, 10, and 21) were further evaluated for senescence characteristics using three different indices: beta-cell identity, beta-cell disallowed genes, and beta-cell senescence and aging-related genes. Additionally, a dendrogram analysis was performed to compare each cluster by analyzing the complete scRNA-seq data. Genes with a positive Log2FC and p-value (p<0.05) were counted for each index in a cluster to calculate the Progression of Senescence (P. S.). The total number of beta-cell identity genes with a positive Log2FC and p-value (p<0.05) was subtracted by the sum of beta-cell disallowed and beta-cell senescence genes with a positive Log2FC and p-value (p<0.05). Using this method, early-stage senescence (clusters 20 and 14), mid-stage (clusters 23 and 10), and late-stage (cluster 21) were identified.

### Global Metabolomics Profiling Using LC-QTOF-MS

Freshly collected mouse plasma samples were diluted (85:15, sample:mixture; v/v) with an acetonitrile:methanol:water (3:3:4) mixture (HPLC-grade, Millipore, Bedford, MA, USA) and 10% methanol, respectively. Each sample (5 μL) was loaded onto an Acquity UPLC BEH C18 (1.7 μm, 2.1 mm × 100 mm; Waters Corp., Milford, MA, USA) column and analyzed using an Agilent 6530 QTOF mass spectrometer (Agilent Technologies). The experimental procedure was performed by the Seoul National University Hospital Metabolomics Core Facility and described in detail in previous studies^115^. The average % coefficient of variance of analytes ranged from 15%– 30%, and the concentrations at the limit of detection are listed in the **Table S4**. Quantified measurements were analyzed using the MetaboAnalyst program^116^. All samples were examined simultaneously in a single run to avoid batch effects, and the overall quality of the analysis procedure was monitored using repeat extracts of a pooled plasma sample (Biocrates, AbsoluteIDQ p180 kit).

### Mitochondria Isolation

Mitochondria were immunopurified from MITO-HA-GFP-expressing MIN6 cells (30 million cells per sample) following the Mito-IP protocol^117^. Mitochondria from shEmpty-tGFP and shSLC25A51-tGFP MIN6 cells were isolated using the Mitochondria Isolation Kit for Cultured Cells (Thermo Fisher Scientific).

## Quantification of NAD^+^ and NADH

### LC-MS/MS in Mitochondria Isolated From HA-Tagged MIN6 Cells

For small polar metabolite (e.g., NAD^+^, NADH) separation and data acquisition on mitochondria samples (diluted in a 40:40:20 methanol:acetonitrile:water mixture with 0.5% formic acid), a Vanquish Horizon (Thermo Fisher Scientific, Waltham, MA) ultra-high-pressure liquid chromatography (LC) system coupled to a Thermo Exploris 480 mass spectrometer (MS) was used. For separation, a Waters (Milford, MA) Xbridge BEH Amide (2.5 μm, 2.1 x 150 mm) column fitted with a VanGuard (2.5 μm, 2.1 x 5 mm) column guard (Waters, Milford, MA) was used. The mobile phases were as follows: Phase A: 95% water/5% acetonitrile and Phase B: 20% water/80% acetonitrile with 10 mM ammonium acetate and 10 mM ammonium hydroxide in both phases. The flow rate was held constant at 0.3 mL/min, and the following gradient conditions were used: 0 min, 100% B; 3 min, 100% B; 3.2 min, 90% B; 6.2 min, 90% B; 6.5 min, 80% B; 10.5 min, 80% B; 10.7 min, 70% B; 13.5 min, 70% B; 13.7 min, 45% B; 16 min, 45% B; 16.5 min, 100% B; and 22 min, 100% B. The samples were held at 4°C, the injection volume was 5 μL, the column was held at 25°C, and the run time was 22 min. The separated metabolites were analyzed on the MS in both positive and negative ionization modes in the same run (switching mode) with the Full MS scan detection mode. The mass spectra were acquired using the following parameters: resolution = 120,000, m/z range = 70–1,000, AGC Target: Standard, Maximum Injection Time: Standard, and run time = 22 min. The ElectroSpray Ionization source parameters for both modes were as follows: Ion Transfer Tube 325°C, Vaporizer 300°C, Spray Voltage 3.5 kV (+)/2.5 kV (-), Sheath Gas 50 (arbitrary units), Aux Gas 10 (arbitrary units), Sweep Gas 1 (arbitrary units), and RF Lens 50%.

### Glutamine/Glutamate-Glo, Ammonia, NAD^+^/NADH-Glo measurements in Pancreatic Islet Cells and Cultured MIN6 Cells

For Glutamine/Glutamate-Glo assay (Promega) and Ammonia assay kit (Abcam), cell lysates from mouse pancreatic islet cells were analyzed according to the manufacturer’s instructions. For NAD measurements in MIN6 cells, thirty million HA-Mito-GFP-expressing MIN6 cells (pMXs-EMPTY-HA-Mito-GFP, FLAG-SLC25A51-HA-Mito-GFP, FLAG-SLC25A51-K91A-HA-Mito-GFP) or shNTC-tGFP and shSLC25A51-tGFP cells were collected for each group. Mitochondrial isolation was performed as described above, and then the NAD^+^/NADH-Glo assay (Promega) was conducted according to the manufacturer’s instructions with 50 µl of the 100 µl mitochondrial fraction for each measurement.

### S961 Treatment *In Vivo*

Prior to the insertion of an osmotic pump, mice were evenly matched and randomized into groups considering their baseline body weights and blood glucose concentrations^19,40,64^. Vehicle (PBS) or 20 nmol S961 diluted in PBS was loaded into an Alzet osmotic pump 2001 and implanted subcutaneously on the back of either 10- or 28- week-old mice, with pumps changed weekly for a total of two weeks.

### IPGTT, ITT, and Insulin Measurements

Prior to the IPGTT, C57BL/6N mice were transferred to clean cages with no food for overnight fasting (12 hr). Mice were weighed to calculate the volume of 20% glucose solution required for intraperitoneal injection (10 μL x body weight in g), and random blood glucose levels were measured at 0 min. Before and after 20% glucose injection, blood was collected from the submandibular vein^118^. The collected serum was stored at -80°C to analyze insulin content within 2 weeks (Ultra Mouse Insulin Elisa Kit, Crystal Chem). For the ITT, C57BL/6N mice were transferred to clean cages and fasted for 4–6 hours before the test. To prepare the insulin solution, two stock solutions were required. First stock: Dilute 50 μL Humulin R 100U/mL (3.5 mg/mL) to 4,950 μL of cold, sterile 1X PBS. Gently invert the solution 3–4 times to mix. Second stock: Dilute 1 mL of the first stock in 9 mL of cold, sterile 1x PBS. Again, gently invert solution 3–4 times to mix. Each mouse was weighed to determine the appropriate dose of the insulin solution, which was administered via intraperitoneal injection. Blood glucose levels were measured and recorded at 15, 30, 60, 90, and 120 min after the initial glucose delivery.

### SA-Beta-Gal Staining

The SA-beta-Gal staining procedure was followed as described previously^119,120^. Briefly, OCT-embedded, snap-frozen, unfixed tissues were cryosectioned at a thickness of 10 µm and stored at -80°C for 24 hours. All tissues were processed and stained within 2–3 days after sacrifice. For SA-beta-Gal staining of frozen tissue sections, pancreatic sections were thawed, fixed, and stained using the Senescence Beta-Galactosidase Staining Kit (Cell Signaling, #9860). For live cell SA-beta-Gal staining using FACS, we used the CellEvent™ Senescence Green Flow Cytometry Assay Kit (Thermo, #C10840) following the manufacturer’s protocol.

### Primary Cell Isolation (Islet Beta Cell, Splenocyte, and Hepatocyte Isolation)

Mice were euthanized, and freshly prepared collagenase P (Roche, Indianapolis, IN) solution (0.5 mg/mL) was injected into the pancreas via the common bile duct. The perfused pancreas was digested at 37°C for 15 min, and the islets were handpicked under a stereoscopic microscope. Cells were then dispersed into a single-cell suspension. Furthermore, pancreatic islet cells were gated, sorted, and purified according to forward and side scatter, with the percentage of beta cells by insulin staining ranging from 85%∼90% (**Figure S1B**), as previously reported^61,62^. For splenocyte and hepatocyte isolation, we conducted the process as previously reported^118^.

### IF Analyses on Pancreatic Tissue Samples

C57BL/6N mice were sacrificed, and pancreases were obtained and preserved in OCT compound for frozen cryotome sectioning. Pancreases embedded in a frozen block were prepared in 5 µm cross-sections for IF staining. Sections were blocked with 5% goat serum (Jackson ImmunoResearch, 005-000-121) in antibody diluent (Dako) for 30 min and incubated overnight at 4°C with the following antibodies: Insulin, Glucagon, or SLC25A51 (Thermo, BS24079R), Cox4 (Proteintech, 66110-1-1g), and p16^INK4A^ (Abcam, Ab54210, Ab211542). After washing with TBST, slides were incubated for 1 hour with the appropriate fluorochrome-attached secondary antibody (Jackson ImmunoResearch). Slides were rinsed and analyzed after mounting with a cover glass. Images were acquired using a confocal microscope (TCS-SP8, Leica, magnification 20x). For quantification, islet images were captured systematically, covering the whole section in confocal mode to evaluate every cluster of insulin-stained cells (3–7 cells) or islets (8 or more cells)/section for SLC25A51 intensity (AU); sections were coded and read blindly. For better microscopic resolution and color contrast, image channels were digitally merged, magnified, and assigned artificial colors using commercially available software programs.

### Flow Cytometry, Intracellular Staining, or Sorting in Tissue-Derived Primary Cells

Isolated mouse beta cells, preadipocytes, splenocytes, or hepatocytes were dispersed using EDTA-PBS for 15 min at 37°C and were resuspended in FACS buffer (0.5% BSA (Millipore) in PBS). Using a FACS Canto (BD), pancreatic islet cells were gated according to forward scatter, with the percentage of beta cells by insulin staining ranging from 85%∼90%. For detecting preadipocytes, splenocytes, and hepatocytes, we followed the gating strategy described in previous reports^118^. For the evaluation of beta-Gal activity, a fluorescent substrate from the flow cytometry assay kit (Thermo, C10841) was used following the manufacturer’s protocol. For intracellular cytokine staining, cells were stained with or without surface markers (e.g., CD45) and then fixed and permeabilized using the Intracellular Fixation & Permeabilization Buffer Set (eBioscience). After permeabilization, an antibody cocktail (IL-1β, CXCL10, IL-6, Insulin) with different fluorochromes was added for intracellular staining. Cells were washed and analyzed in FACS buffer. Data were analyzed using FlowJo software (Treestar, Ashland).

### SASP Analysis in Mouse Serum Using the Multiplex-Cytokine Assay

Mouse serum obtained from chronologically aged mice (12W, 30W, 90W) and treatment groups (12W-C, 30W-C, 30W-S) was stained and analyzed using a multiplexed bead-based assay for mouse proinflammatory chemokines (Legendplex, 740446, BioLegend).

### Cell Lines, Plasmids, and induction of senescence

MIN6 cells were cultured in DMEM supplemented with 20% fetal bovine serum (FBS). HEK293FT cells for virus production were grown in DMEM supplemented with 10% FBS. Cells were incubated in a humidity-controlled environment at 37°C with 5% CO_2_ (Thermo Fisher Scientific Forma 371) and were used up to 10 passages. pGenLenti-empty, pGenLenti-CDKN1A, pGenLenti-CDKN2A, and pGenLenti-NFE2L2 were obtained from Genscript. pMXs-FLAG-SLC25A51, pMXs-FLAG-SLC25A51-K91A mutant, and pMXs-HA-GFP vectors are from a previous study^50^. pMXs-FLAG-SLC25A38 and pMXs-SOD2-CAT-GFP vectors were generated in this study. pLKO.1-tGFP-shNTC (non-targeting control, SHC016) and pLKO.1-tGFP-shSLC25A51 were purchased from Mission shRNA Custom DNA (Millipore Sigma). Senescence was induced by 10 Gy ionizing radiation using RS-2000 (Rad Source Technologies) or treating cells with H_2_O_2_ (200 μM).

### Virus Production in HEK293FT Cells

HEK293FT cells were co-transfected with the VSVG envelope plasmid, retroviral packaging plasmids Gag-Pol, and pMXs, pLKO.1, or pLenti plasmids. The plasmids included pMXs-HA-MITO-GFP (Addgene, #83356), pMXs-FLAG-SLC25A51 (Addgene, #133247), pMXs-FLAG-SLC25A51-K91A (Addgene, #133248), pMXs-FLAG-SLC25A38 (Addgene, #218412), pMXs empty vectors, pLKO.1-Neo-CMV-tGFP-shNTC (non-targeting control; Mission shRNA, Millipore, SH016), pLKO.1-Neo-CMV-tGFP-shSLC25A51 (Mission shRNA, Millipore), pLenti-Nfe2l2, pLenti-Cdkn2a, or pLenti-Cdkn1a (Genscript). Transfections were performed using 2.0 x 10^6^ cells in 6-well plates in RPMI containing polybrene (10 μg/mL; Millipore). The culture medium was changed 8 hours after transfection. The virus-containing supernatant was collected 48 hours after transfection, spun, and filtered to eliminate debris and cells. Subsequently, virus-containing supernatants were used to generate MIN6 stable cell lines.

### Generation of MIN6 Stable Cell Lines

MIN6 cells were transfected with different viruses collected from HEK293FT cultured media, as described above. After an 18-hour incubation with the virus, the medium was changed to fresh culture medium containing the appropriate selection reagent (pMXs: blasticidin, pLenti: puromycin, pLKO.1: neomycin) and selected for 72 hours. MIN6 cells with HA-GFP or tGFP were sorted using a FACS Aria sorter (BD).

### MIN6 Cell Sorting

For sorting MIN6 stable cell lines, GFP-positive (pMXs-HA-GFP or pLKO.1-NEO-CMV-tGFP-shEMPTY or pLKO.1-NEO-CMV-tGFP-shSLC25A51) cells were acquired and sorted using a FACS Aria cell sorter (BD). For evaluating mitochondrial morphology and function, TMRE, MitoTracker, and MitoSOX were used to stain either MIN6 or pancreatic beta cells. Cells were washed and analyzed in FACS buffer. Data were analyzed using FlowJo software (Treestar, Ashland).

### mRNA Quantification by qPCR

Total RNA was purified using the Rneasy Mini Kit (Qiagen) as per the manufacturer’s protocol. cDNA was synthesized by reverse transcription of 1 μg of total RNA using AMV Reverse Transcriptase (Promega) following the manufacturer’s instructions. Quantitative real-time PCR was performed in a 20 μL reaction mixture containing 1 μL of cDNA, 10 pM of each primer, and 10 μL of SYBR Select Master Mix (Thermo Fisher Scientific) using an Applied Biosystems 7500 Real-Time PCR System (Thermo Fisher Scientific). Specific primers used for qPCR are listed in **Table S5**.

### Western Blot Analysis

Cell lysates were subjected to electrophoresis using 4–20% precast gels (Thermo Fisher Scientific). The resolved gels were transferred to PVDF membranes, blocked with 5% skimmed milk in Tris-buffered saline containing 0.05% Tween-20 (TBS-T), and incubated with primary antibodies against SLC25A51 (Mouse, Thermo BS-24079R), SLC25A38 (Abcam, ab133614), IGF1R (Cell Signaling, 9750S), p16^INK4A^ (Novus, NBP2-37736), p21^cip1/waf1^ (Invitrogen 14-6715-81, Abcam ab188224), p53 (Genetex, GTX102965), γ-H2AX (Cell Signaling, 9718S), NRF2 (Proteintech, 16396-1-AP), NNT (Proteintech, 13442-2-AP), NAMPT (Proteintech, 66385-1-1g), TOM20 (Proteintech, 11802-1-AP), Citrate Synthase (Cell Signaling, 14039S), β-actin conjugated with HRP (Cell Signaling, 5125S), CD3e (Invitrogen, MA5-16622), Insulin (Abcam, ab181547), Cyp8b1 (Thermo, PA5-37088), and OXPHOS antibody cocktail (Abcam, ab110413) at 4°C overnight. The membranes were incubated with HRP-conjugated secondary antibodies at room temperature for 2 hours and developed using Pico chemiluminescent substrate (Thermo Fisher Scientific) and imaged using an Amersham Imager 600 (GE Healthcare). At least three trials of Western blot analysis were performed.

## Bioinformatics and Statistical Analyses

### scRNA-seq Gene Expression Analysis

Single-cell gene expression data were analyzed using Cell Ranger v5.0.0 (10x Genomics; https://support.10xgenomics.com/single-cell-gene-expression/software/pipelines/latest/what-is-cell-ranger). Briefly, raw BCL files from the Illumina HiSeq platform were demultiplexed to generate FASTQ files using ‘cellranger mkfastq’. Then, these raw FASTQ files were analyzed using ‘cellranger count’. The ‘cellranger count’ step includes mapping to the *Mus musculus* reference genome (mm10-2020-A), measuring gene expression with unique molecular identifiers (UMIs) and cell barcodes, determining cell clusters, and conducting differential gene expression analysis. The ‘count’ function can take input from multiple sequencing runs on the same library. The resulting raw count matrices were imported into Seurat 4.3.0. All genes expressed in ≥3 cells and all cells with at least 200 detected genes were used in downstream analysis. Rare subsets of cells with an abnormally high number of genes detected, suggesting potential multiplets, and low-quality or dying cells were filtered out if they had more than 6,000 genes or mitochondrial counts exceeding 15%. After filtering, the final count matrix contained 23,258 genes across 6,977 cells. Gene expression values were scaled and normalized for each gene across all integrated cells. Clustering and UMAP analysis were performed based on the statistically significant principal components. Significant top cluster markers for each cluster compared to all remaining cells were determined using the Wilcoxon rank sum test (min.pct=0.25, logfc.threshold=0.25, only the positive ones were filtered). Gene enrichment and functional annotation analysis for significant gene lists from two different results were performed using the g:Profiler tool (https://biit.cs.ut.ee/gprofiler/).

### GO and KEGG Analysis

GO and functional annotation analysis for significant probe lists were performed using KEGG (www.genome.jp/kegg/)^121^ (**Table S1**). All data analysis and visualization of differentially expressed genes was conducted using R 3.1.2 (www.r-project.org).

### Network, STRING, Hierarchical Clustering, and Heatmap Analysis

Metabolomics data were further analyzed for network and pathway enrichment analyses were performed using MetaboAnalyst (version 5.0) (https://www.metaboanalyst.ca/). STRING analysis was performed using STRING (https://string-db.org/). Hierarchical clustering was conducted using the Ward variance minimization algorithm (method=’Ward’) with Euclidean distance (metric=’euclidean’) in SciPy (1.13.0) (**Figures 3C, S2A**). For heatmap analysis, the colors of individual cells in the heatmaps represent the Log2 fold changes obtained using the 10x Genomics Loupe Browser v8.0.0 (**Figures 3A, S2A**). The values represent the ratio of normalized mean gene UMI counts of individual clusters compared to all other clusters.

Genes with a smaller p-value are considered differentially expressed in the heatmap (**Figures 3A, S2A**). P-values were adjusted using the Benjamini–Hochberg correction for multiple feature tests. Hierarchical clustering was performed using SciPy, applying the Ward variance minimization algorithm with Euclidean distance (**Figures 3A, S2A**).

### Cell Type Clustering Using Seurat

Calculation of single-cell expression matrices by Cell Ranger was performed using Seurat package for filtering, data normalization, dimensionality reduction, clustering, and gene differential expression analysis^122^. The filtered scRNA-seq data were analyzed as follows:

1. Low-quality, multiplets, and mitochondrial contamination check: Cells with fewer than 200 genes, more than 8,000 genes, or a mitochondrial gene ratio greater than 25% were excluded. Doublets were detected using three standards: (1) The number of genes (nFeature), representing genes with at least one UMI count detected in each cell, which helps remove empty droplets or low-quality cells and identify doublets or multiplets. (2) The total number of molecules (nCount), representing the number of UMIs detected within a cell. (3) The percentage of reads mapped to the mitochondrial genome.
2. Aggregation for multiple datasets: Batch effects were corrected using anchors and canonical correlation analysis from the Seurat R library when aggregating multiple datasets.
3. Normalization and scaling: A global-scaling normalization method, “LogNormalize,” was applied to normalize gene expression measurements for each cell by total expression, multiply by a scale factor (10,000 by default), and log-transform the result.
4. Detection of variable genes: Highly variable genes showing a clear relationship between variability and average expression were extracted for dimensionality reduction analysis.
5. Scaling and centering data: The percentage of mitochondrial gene content was regressed against each feature, and the resulting residuals were scaled and centered. These residuals were used for dimensionality reduction and clustering analysis.
6. Principal component analysis (PCA): PCA was performed on the scaled data for graph-based clustering, selecting the number of principal components, k: 15.
7. Clustering cells: To find the graph-based clusters, we first constructed a K-nearest neighbor graph based on the Euclidean distance in PCA space, and refined the edge weights between any two cells based on the shared overlap in their local neighborhoods (Jaccard similarity). We also applied modularity optimization techniques such as the Louvain algorithm (default) to iteratively group cells together, with the goal of optimizing the standard modularity function. As UMAP/tSNE can help place cells with similar local neighborhoods in high dimensional space together into a low-dimensional space, we used UMAP for dimensionality reduction. UMAP is an algorithm for dimension reduction based on manifold learning techniques and ideas from topological data analysis^123^.
8. Visualization: UMAP clustering results were visualized using the 10x Genomics Loupe Browser (version 8.0.0). Following this pipeline, we processed scRNA-seq data from 26,662 high-quality pancreatic tissue cells to create a single-cell mouse pancreas atlas. We identified 26 clusters representing senescence and diabetes-associated pancreatic islet cell types.

### Annotation of Cell Types Using CellKB

To confirm the classification of islet cell types, we used a three-step procedure: (1) SingleR, a reference-based scRNA-seq annotation^124^, (2) comparison of cell numbers between groups, and (3) comparison of senescent beta-cell genes in each cluster. CellKB^Pro^ (version 2.1.2) (http://www.cellkb.com) is a knowledgebase of author-defined cell type marker gene sets (n=33,064) that can be rapidly searched for matching cell types (n=2,407), as previously reported ^125^. CellKB^Pro^ collects marker gene sets directly from publications describing primarily scRNA-seq and selected bulk RNA-seq or microarray experiments. It contains extensive cell type, tissue, and disease annotations for each cell type as described in the publication. To obtain a detailed and reliable pancreatic islet cell classification, we uploaded DEGs of each cluster (in descending order of fold change). The query genes of each cluster were compared with every marker gene set in the database, and the best match was identified using a match score. The rank score was calculated based on the number of common genes between the query and the cell type, their ranks, their rank differences, and the total number of significant genes in the cell type. Higher match scores were assigned to cell types sharing highly ranked genes with the query. CellKB^Pro^ used an exponential function to score the matching genes, emphasizing higher-ranked matches more than lower-ranked ones. The algorithm also accounted for differences in gene list sizes between the query and various cell types to prevent disregarding cell types with fewer marker genes. After uploading the data, we obtained a list of relevant cell types (up to 100, ranked from highest to lowest score) for the DEGs of all 26 clusters (**Figure 1H**). According to the rank-based score, 10 clusters (0, 2, 5, 10, 11, 12, 14, 20, 21, and 23) were classified as beta-cell clusters. Furthermore, among these beta-cell clusters, clusters 10, 11, 12, 14, 20, 21, and 23 exhibited a “senescent metaplastic cell” type (**Table S2**). Thus, senescent beta-cell clusters were identified as 10, 14, 20, 21, and 23. Before confirming the expression levels of senescent genes in beta-cell clusters (**Figure 3A**), we compared cell numbers between groups in each cluster (**Figures 2K, S2E**). Clusters 10, 14, 20, 21, and 23 showed a consistent increase in cell numbers, while clusters 2, 5, 11, and 12 showed a lower cell proportion (%) in 30W-S compared to 12W-C or 30W-C groups (**Figures 2K, S2E**). To reconfirm whether clusters filtered by steps 1 and 2 were indeed senescent beta-cell clusters, we compared gene expression in each beta-cell cluster with senescent beta-cell gene sets from previous studies characterizing senescent beta cells, using 69 genes (22 genes for hallmark beta-cell identity genes^19,70,126,127^, 21 genes for beta-cell senescence^19,22,27,39,60,63,93,128,129^, and 26 genes for beta-cell disallowed genes^71,72^):

A. Beta-cell identity genes: *Rtx6, Pax6, Abcc8, Gck, Syt13, Vdr, Nkx2-2, Chga, Neurod1, Kcnj11, Ins1, Ins2, Mafa, Iapp, Nkx6-1, Pdx1, Dcx, Foxa2, Pklr, Elp4* (*Sst* and *Lmo2* as negative controls) ^19,70,126,127^,
B. Beta-cell senescence genes: *Ulk3, Arntl, Igf1r, Gls, Mapk14, Cdkn1a, Trp53, Map2k1, Sirt1, Id2, Il6, Cdkn2a, Il1a, Prkcd, Bambi, Plaur, Gls2, Cav1, Tbx2, _Tbx3, Il1b_* 19,22,27,39,60,63,93,128,129.
C. Beta-cell disallowed genes: *Fcgrt, Zcchc24, Arhgdib, Igf2bp2, Cox5a, Uqcrq, Zfp36l1, Pld2, Smad3, Cat, Selenbp1, Parp3, Zyx, Klhl5, Ldha, Arap2, Lmo4, Slc16a1, Tgm2, Yap1, Ndrg2, Mylk, Hpgds, Rnf144a, MgII, Rarres2* ^71,72^.

These genes were assessed using Log2 fold changes (**Figures 3A**). We confirmed that clusters 10, 14, 20, 21, and 23 are senescent beta-cell clusters.

### Analysis of Gene Expression in Publicly Available Datasets

Beta-cell RNA-seq data were obtained from human male and female datasets (E-MTAB-5061, GSE81608)^104,130^. SLC25A family genes, beta-cell identity genes, beta-cell disallowed genes, and beta-cell senescence genes were analyzed in each dataset. Additional pancreatic islet cell datasets (GSE73727^131^, GSE85241^132^, GSE87375^105^) were analyzed for SLC25A51 mRNA levels. Non-pancreatic islet cell types were also analyzed using aging datasets from *A. thaliana* and *S. cerevisiae* (GSE43616^133^, GSE210032^134^) and human cell lines (WI38: GSE130727^54^; MSC, IMR90 HUVEC: GSE98440^56^; HDF161: GSE93535^59^; RPE-1: GSE83647^57^; LS8817:

GSE74620^58^; MRC5: GSE64553^55^). The data were visualized using heatmaps or dot-plot graphs in GraphPad Prism.

## Quantification and Statistical Analysis

Data are presented as mean ± standard error of the mean (SEM). GraphPad Prism was used for two-tailed t-tests, one-way, or two-way ANOVA to assess significance between groups, followed by Bonferroni’s multiple comparison test.

## Declaration of interests

N.K. is an inventor on a patent application filed by the Whitehead Institute for Biomedical Research relating to work described in this paper. Y.M.C. is an outside director of Daewoong Pharmaceutical and received a consultation fee from LG Chemical. All other authors declare that they have no competing interests.

## Author Contributions

BSK and NK conceptualized the experiments, interpreted data, and prepared the figures. BSK was responsible for plasmid and viral production to generate stable cell lines and performed statistical analyses. BS Kong and SR Hong conducted beta-cell experiments. BSK, EGRH, SRH, SYS, and CC performed mouse experiments. BSK, NK, and YMC conceived the project, acquired funding, and BSK and NK wrote the manuscript. All authors reviewed and approved the manuscript.

## Acknowledgements

We thank Dr. Dajung Kim (Metabolomics Core Facility, Department of Transdisciplinary Research and Collaboration, Biomedical Research Institute, Seoul National University Hospital) for LC/MS-MS metabolomics analyses. We also thank Aparna Vijayakumar and Dr. Lan Hai for plasmid and virus production, as well as animal husbandry. We thank Drs. Kristopher Sarosiek and Atomu Yamaguchi for assistance with irradiation experiments. This research was supported by a William F. Milton Fund grant and a Dean’s Fund for Scientific Advancement Incubation Award to NK, the Basic Science Research Program through the National Research Foundation of Korea (NRF), funded by the Korean government (MSIT) (NRF-2022R1C1C2003297) and the Seoul National University Hospital Research Fund (042022-4080) to BSK. Additional support was provided by an R35/MIRA (GM151097), a Damon-Runyon Rachleff Innovation Award (73-22), and JDRF (1-INO-2023-1342-A-N). The Harvard College Program for Research in Science and Engineering (PRISE) provided support to CC. We thank all members of the Kory lab, Dr. Nika Danial, Dr. Cristina Aguayo-Mazzucato, and Dr. Christopher Whiley for their helpful comments, and Dr. Julie Gosse for editing the manuscript.

## Supplementary Figures

**Figure S1.**
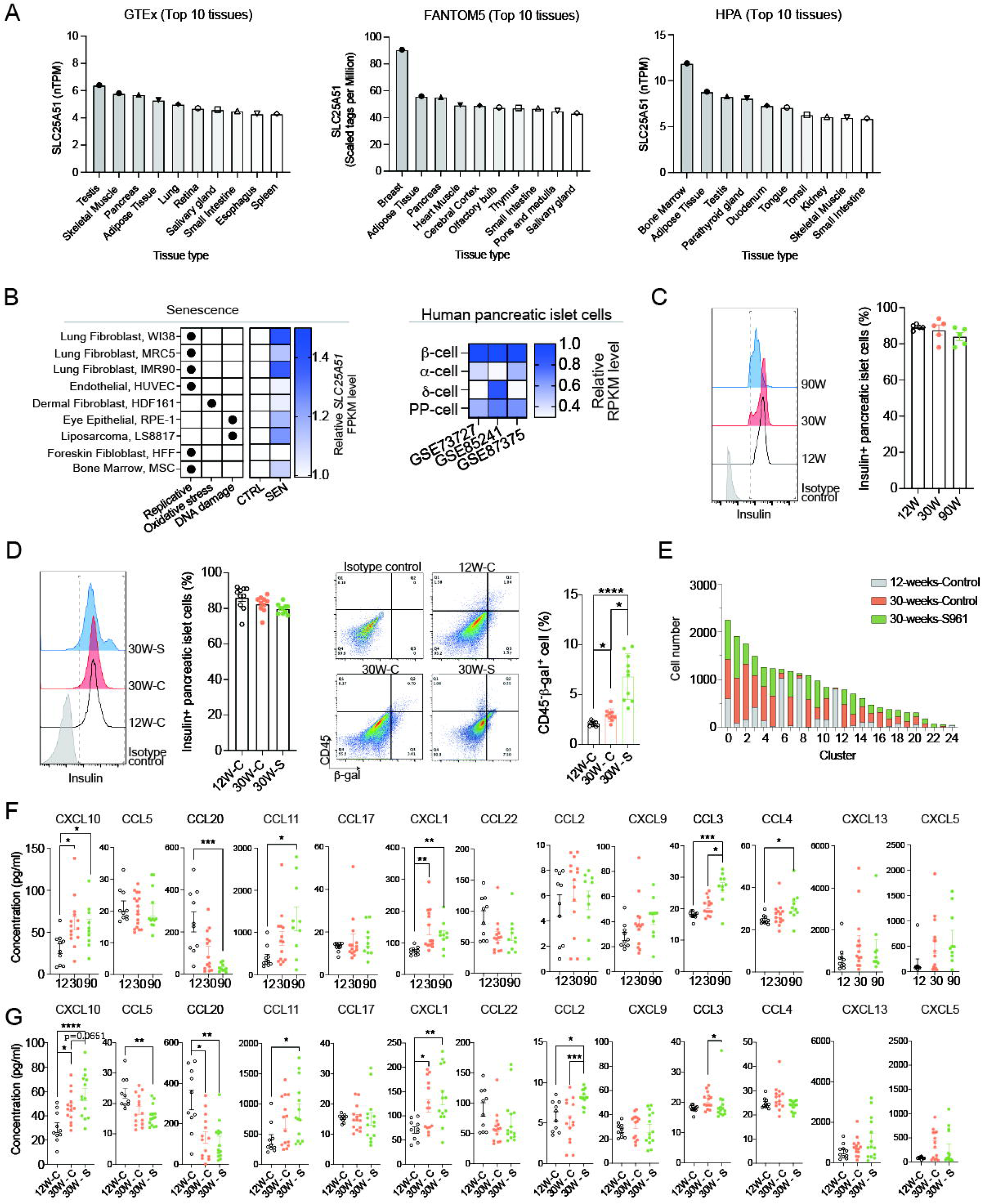
(**A**) Human *Slc25a51* mRNA levels were compared between different tissues from publicly available datasets (GTEx, Fantom5, and HPA). (**B**) Human cell lines before and after induction of senescence by extensive replication, oxidative stress, or DNA damage were analyzed for *Slc25a51* mRNA levels (WI38: GSE130727; MSC, IMR90, HUVEC: GSE98440; HDF161: GSE93535; RPE-1: GSE83647; LS8817: GSE74620; MRC5: GSE64553). Human pancreatic islet cells (alpha, beta, PP, delta-cells) scRNA-seq data (GSE73727, GSE85241, GSE87375) were analyzed for SLC25A51 in RPKM levels (**Table S3**). (**C, D**) Beta ce€ isolated from (**C**) 12W, 30W, and 90W (n=5/group) or (**D**) 12W-C, 30W-C, and 30W-S groups (n=10/group) were stained with (**C, D**) insulin, (**D**) CD45, and SA-beta-gal. SA-Beta-gal frequency in pancreatic islets was analyzed in insulin^+^CD45^-^SA-beta-gal^+^ cells from 10 mice per group. One-way ANOVA; error bars represent SEM. *p < 0.05, ****p < 0.0001. (**E**) Cell numbers of every cluster from scRNA-seq (Figure 1) were compared between groups (12W-C, red; 30W-C, blue; 30W-S, green). (**F, G**) Senescence-associated secretory phenotype (SASP) was analyzed in mouse serum (n=10/group) using the Legendplex mouse proinflammatory chemokine panel. Heatmap and individual dot plots are assessed for (**G, I**) 12-, 30-, 90-week-old mice and for (**H, J**) 12W-C, 30W-C, and 30W-S groups; One-way ANOVA; error bars represent SEM. *p < 0.05, **p < 0.01, ***p < 0.001.

**Figure S2.**
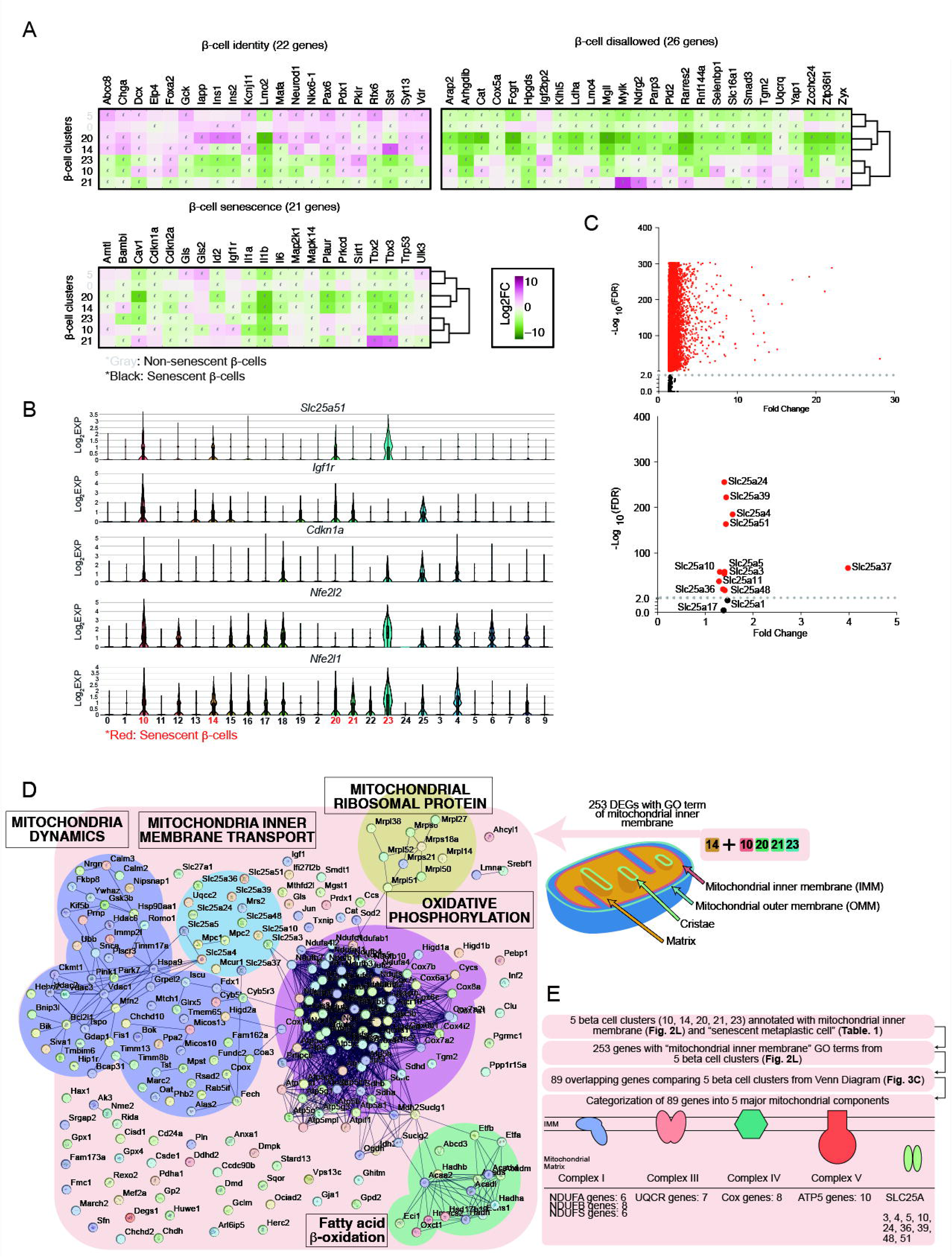
Beta-cell identity genes, beta-cell disallowed genes, beta-cell senescence, and aging-related genes were analyzed in clusters 0, 5, 10, 14, 20, 21, and 23. (**A**) *Slc25a51*, *Igf1r*, *Cdkn1a*, *Nfe2l2*, and *Nfe2l1* genes were compared in all 26 clusters. Expression of genes is in Log2(Ex€ssion) (Log2EXP). (**B**) Volcano plots for all 11,668 DEGs (up) and SLC25A family genes (bottom). The dotted line indicates FDR<0.05. (**C**) A total of 253 genes showed the GO term “mitochondrial inner membrane” in clusters 10, 14, 20, 21, and 23. With these 253 genes, we performed STRING analysis indicating that these genes were involved in “mitochondrial inner membrane transport,” “mitochondria dynamics,” “mitochondrial ribosomal protein,” “oxidative phosphorylation,” and “fatt€cid beta-oxidation.” (**E**) Diagram showing the unbiased approach to mine mitochondrial-related genes significant inescence.

**Figure S3.**
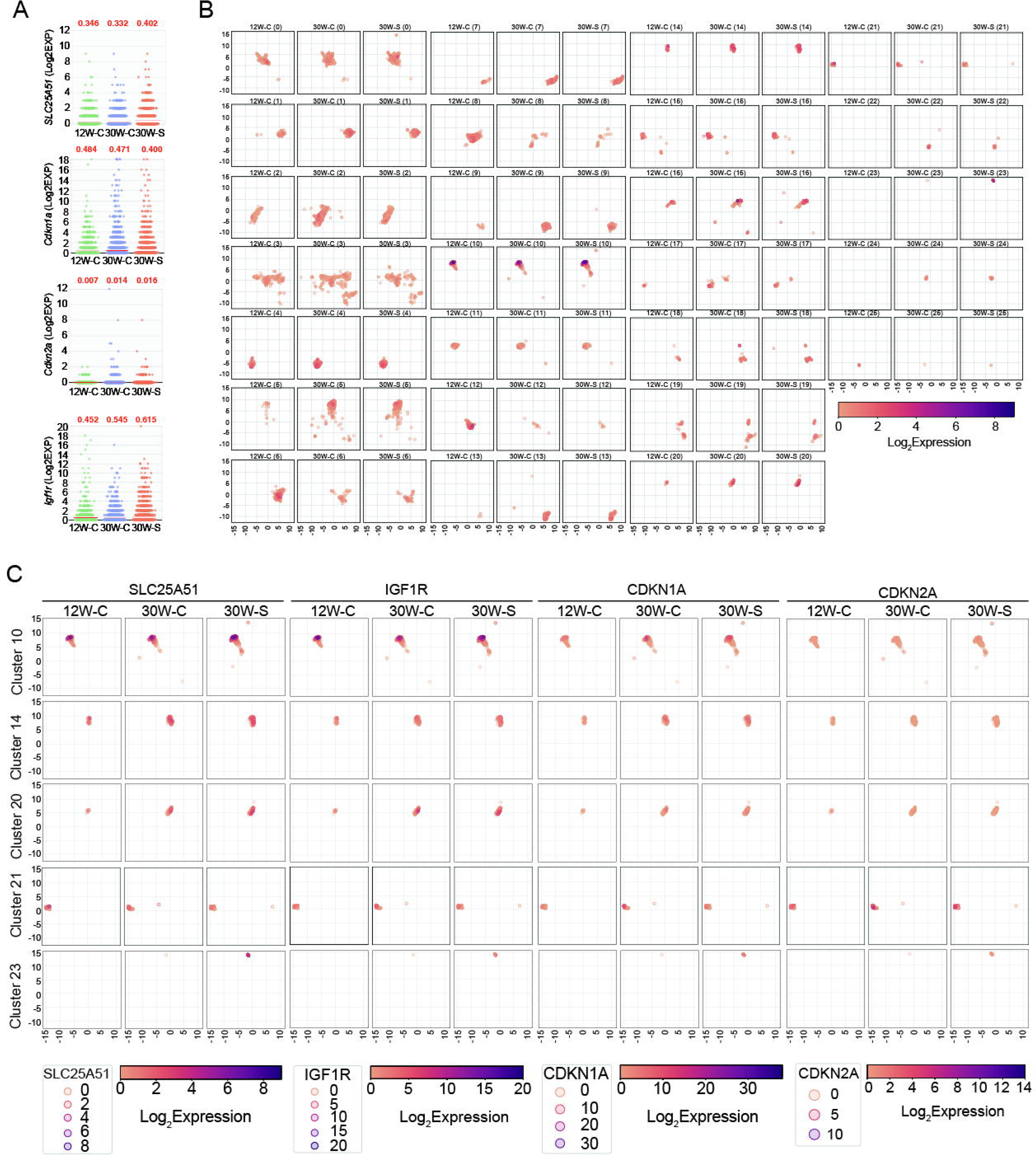
(**A**) *Slc25a51*, *Cdkn1a*, *Cdkn2a*, and *Igf1r* expression levels (Log2EXP) per cell were analyzed in each group (12W-C, 30W-C, 30W-S; n=21,853 cells). Average Log2EXP expression values are in red for each gene and each group. (**B, C**) *Slc25a51* expression levels in different groups (12W-C, 30W-C, 30W-S) and clusters (0–25 clusters) were compared in Log2EXP.

**Figure S4.**
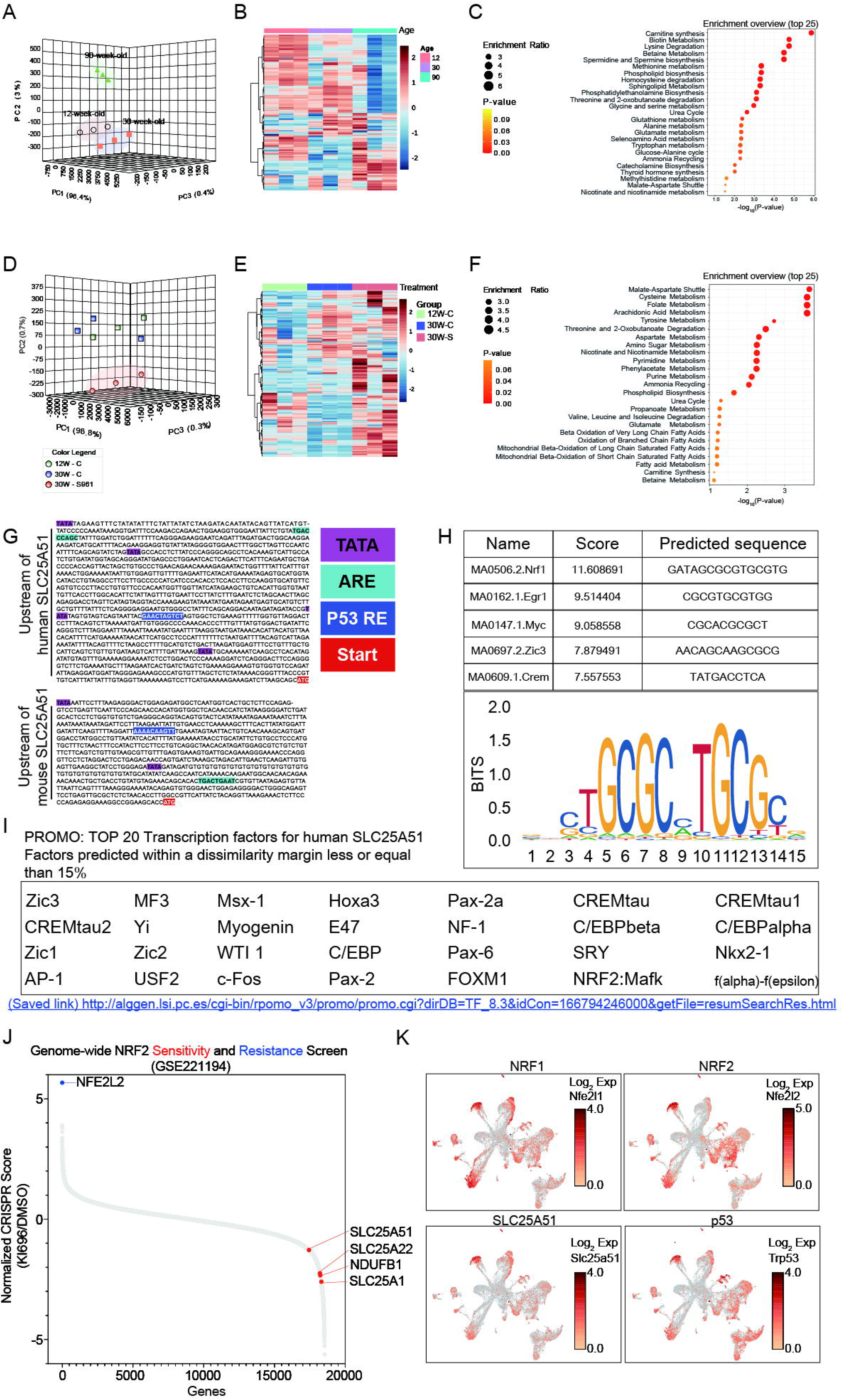
To obtain an overview of systemic metabolic changes in (**A-C**) chronologically aged or (**D-F**) S961-treated mice, we performed targeted metabolomics on mouse serum (n=3/group). Serum samples were collected from 12-week (12W), 30-week (30W), and 90-week (90W) mice as well as from 12W-C, 30W-C, and 30W-S groups for metabolomics analysis. (**A, D**) Principal component analysis (PCA) revealed clear shifts in metabolite profiles among 12W, 30W, and 90W mice, as well as between 12W-C, 30W-C, and 30W-S groups driven by multiple components. (**B, E**) Using the MetaboAnalyst database, we performed hierarchical €stering between groups. (**C**) 30W and 90W or (**F**) 30W-C and 30W-S were compared to analyze the overview of enriched metabolism pathways. (G) Mouse and human *Slc25a51* upstream sequence showing antioxidant-response element (ARE), p53 response element (p53 RE), TATA box, start codon for *Slc25a51*, and arrow indicating the *Slc25a51* gene sequence, and (**H**) Predicted top 5 transcription factors for mouse *Slc25a51* enhancer (top) and bona fide *Slc25a51* target genes (bottom) analyzed by JASPAR (http://jaspar.genereg.net/). (**I**) Top 20 transcription factors anal<u>yzed using th</u>e PROMO program (http://alggen.lsi.upc.es/cgi-bin/promo_v3/promo/promoinit.cgi?dirDB=TF_8.3) for the human *Slc25a51* gene. (J) To search for genes that may predict sensitivity or resistance to NRF2 activation, Weiss-Sadan et al. performed a genome-wide CRISPR screen in the KEAP1-dependent cell line CALU6. Following infection with the sgRNA library, cells were grown for 11 population doublings in the presence of 1 μM of KI696 or vehicle control. As validation, NRF2 appeared as the to-scoring gene that mediated resistance when depleted in KEAP1-1-dependent CALU6 cells. Among many oxidative phosphorylation-related genes, *Slc25a51*, Slc25a22, *Ndufb1*, and Slc25a1 were enriched in KEAP1-dependent CALU6 cells. (K) *Slc25a51*, *Trp53* (p53) *Nfe2l1* (Nrf1), and *Nfe2l2* (Nrf2) expression levels were analyzed by UMAP in pancreatic islet scRNA-seq from Figure 2.

**Figure S5.**
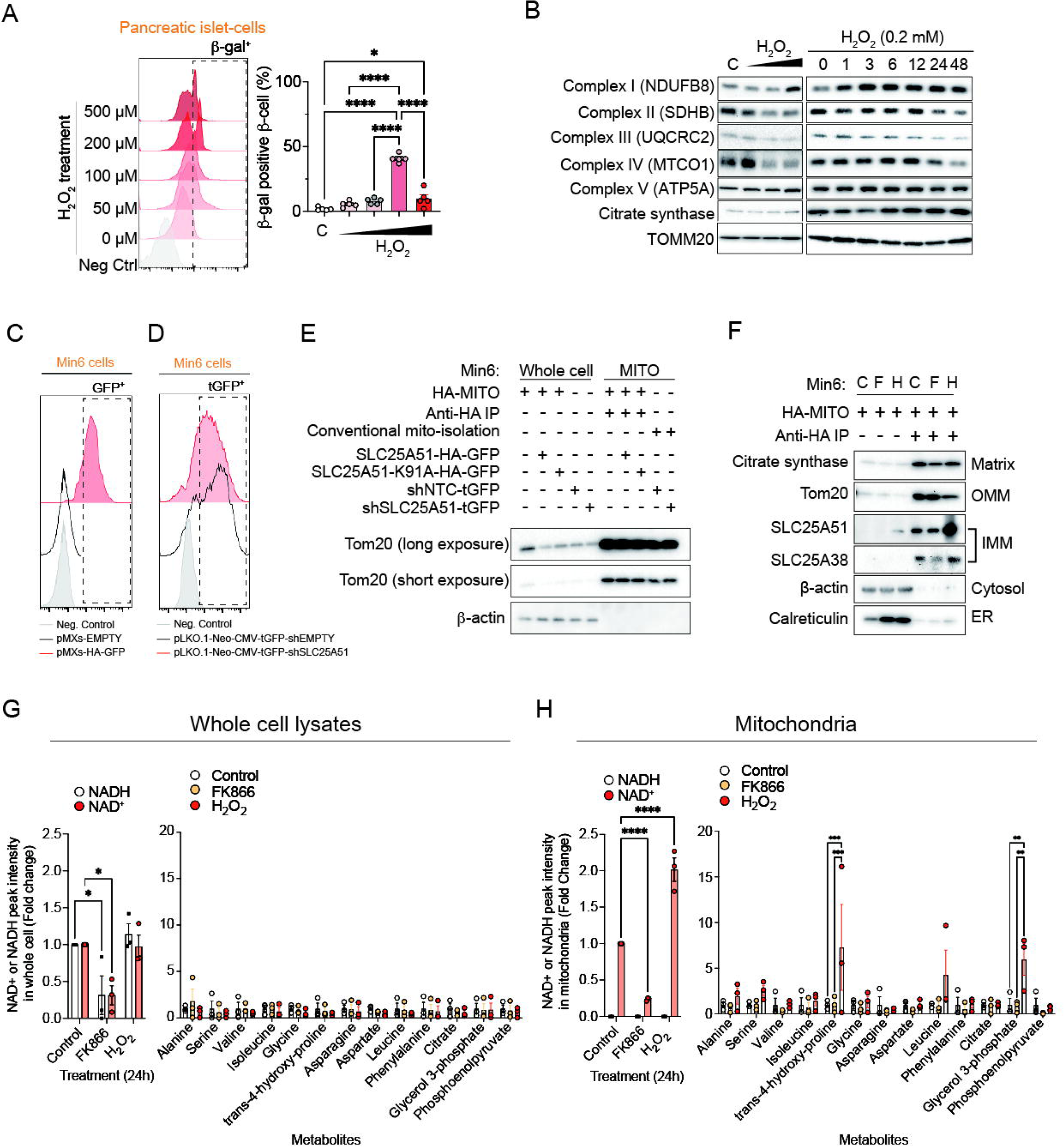
Pancreatic islet cells were isolated from 30-week-old C57BL/6N male mice (n=5/group). The cells were treated with H_2_O_2_ and stained for SA-beta-Gal activity; Two-way ANOVA, error bars represent SEM, *p<0.05, ***p<0.001, ****p<0.0001. (A) MIN6 cells were treated with H_2_O_2_ in a time- and concentration-dependent manner to probe for mitochondrial ETC complexes. (**C-E**) MIN6 cells were transfected with le€viral vectors to generate: (**C**) pMXs-Empty and pMXs-HA-GFP; (**D**) pLKO.1-shEMPTY and pLKO.1-shSLC25A51 MIN6 cells. Each stable cell line was assessed for GFP or tG€signals using flow cytometry. (**E**) Mitochondria were isolated from different stable cell lines and then probed for Tom20 and beta-actin. (**F-H**) pMXs-HA-Mito-GFP-transfected MIN6 cells were treated with PBS, FK866 (2 nM, 24 hours), or H_2_O_2_ (200 μM, 24 hours). Cells were processed to isolate whole-cell lysates and purified mitochondria by MITO-IP. (**F**) Samples were probed for mitochondrial matrix protein (citrate synthase), mitochondrial OMM protein (Tom20), mitochondrial IMM proteins (SLC25A51, SLC25A38), cytosolic protein (beta-actin), and ER protein (calreticulin). Furthermore, metabolites were extracted from (**G**) whole-cell lysates and (**H**) purified mitochondria to perform LC-MS analysis (n=3/group). Two-way ANOVA, error bars represent SEM, *p<0.05, **p<0.01, *0.001, ****p<0.0001.

**Figure S6.**
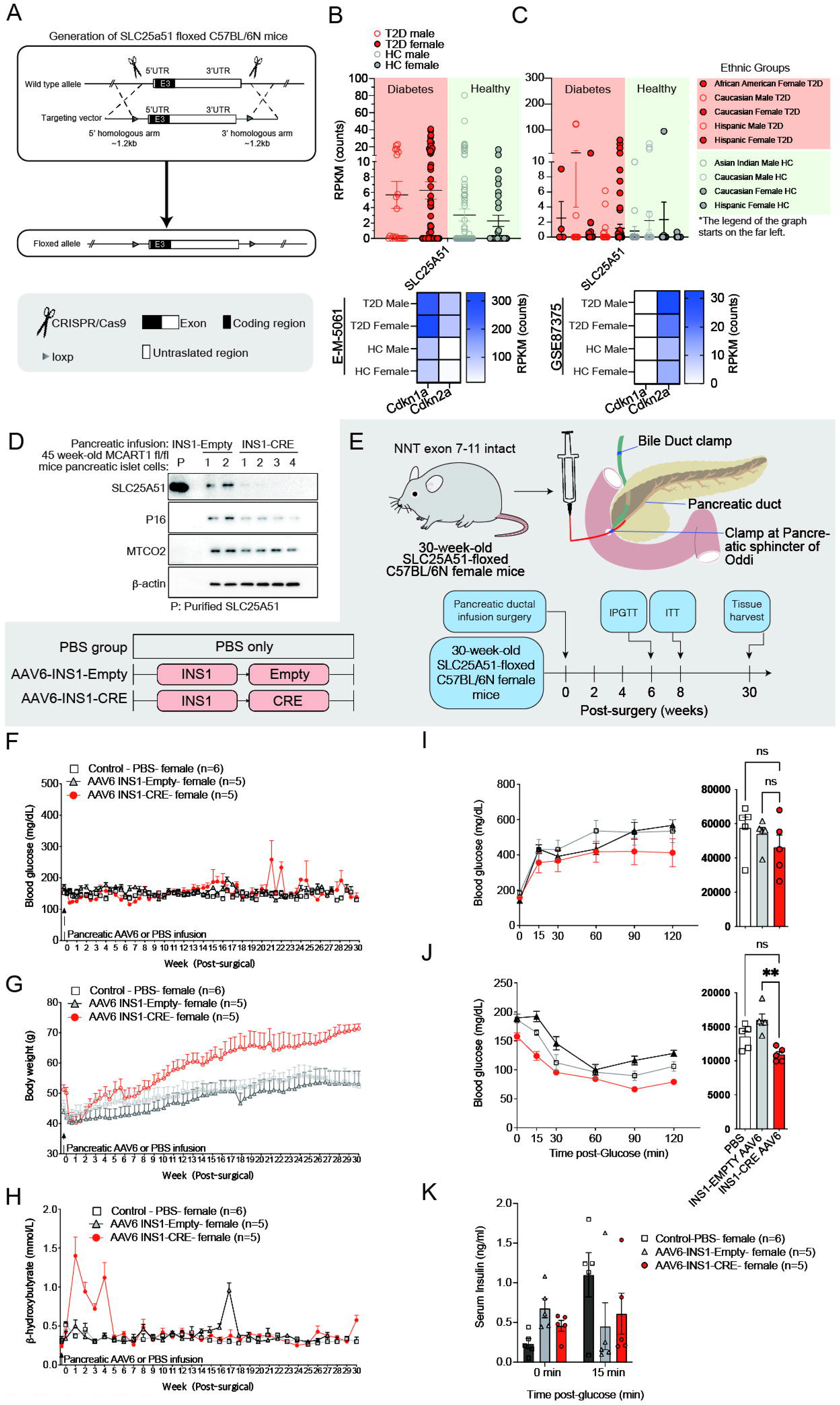
**(A)** (**A**) A diagram depicting the generation of SLC25A51-floxed C57BL/6N mice using a CRISPR-Cas9-based approach by Biocytogen Pharmaceuticals. (**B, C**) SLC25A51 expression per cell was compared between T2D patients and healthy controls using two different human beta-cell scRNA-seq datasets (E-MTAB-5061, GSE81608) (**Table S3**). Heatmaps of *Cdkn1a* (p21) and *Cdkn2a* (p16) expression were also analyzed in both datasets. (**D**) 30-week-old SLC25A51-floxed C57BL/6N male mice were infused with either AAV6-INS1-Empty or AAV6-INS1-CRE viruses via pancreatic ductal infusion. Mice were sacrificed 15 weeks post-surgery. SLC25A51, p16^INK4A^, MTCO2, and beta-actin protein expression levels were analyzed in pancreatic beta cells isolated from 45-week-old INS1-EMPTY (n=2) and INS1-CRE (n=4) groups. (**E-K**) Pancreatic ductal infusion of either PBS, AAV6-INS1-Empty, or AAV6-INS1-CRE was performed in each group of female C57BL/6N mice (n=5–6 per group). (**F**) Random blood glucose levels and (**G**) body weight were measured twice a week, and (**H**) beta-hydroxybutyrate levels were measured once a week for each mouse. (**I-K**) Six or eight weeks post-surgery, female mice underwent an IPGTT and ITT. Graphs show blood glucose levels at (**I, J**) each time point and area under the curve. (**I**) During the IPGTT, serum was collected from each group at 0 and 15 min to measure (**K**) serum insulin levels. Two weeks after the IPGTT, (**J**) an ITT was performed to test insulin sensitivity. Two-way ANOVA; error bars represent SEM; **p < 0.01.

## Notes

### Competing Interest Statement

Byungsoo Kong, Nora Kory, and Young Min Cho are inventors on a provisional patent application filed by Harvard University. Young Min Cho is an outside director of Daewoong Pharmaceutical and received a consultation fee from LG Chemical.

### Summary of Updates

Author list and funding information was edited.

## References

1. Campisi, J. (2013). Aging, cellular senescence, and cancer. Annu Rev Physiol 75, 685–705. 10.1146/annurev-physiol-030212-183653.

2. McReynolds, M.R., Chellappa, K., Chiles, E., Jankowski, C., Shen, Y., Chen, L., Descamps, H.C., Mukherjee, S., Bhat, Y.R., Lingala, S.R., et al. (2021). NAD(+) flux is maintained in aged mice despite lower tissue concentrations. Cell Syst 12, 1160–1172 e1164. 10.1016/j.cels.2021.09.001.

3. Camacho-Pereira, J., Tarrago, M.G., Chini, C.C.S., Nin, V., Escande, C., Warner, G.M., Puranik, A.S., Schoon, R.A., Reid, J.M., Galina, A., and Chini, E.N. (2016). CD38 Dictates Age-Related NAD Decline and Mitochondrial Dysfunction through an SIRT3-Dependent Mechanism. Cell Metab 23, 1127–1139. 10.1016/j.cmet.2016.05.006.

4. Cercillieux, A., Ratajczak, J., Joffraud, M., Sanchez-Garcia, J.L., Jacot, G., Zollinger, A., Metairon, S., Giroud-Gerbetant, J., Rumpler, M., Ciarlo, E., et al. (2022). Nicotinamide riboside kinase 1 protects against diet and age-induced pancreatic beta-cell failure. Mol Metab 66, 101605. 10.1016/j.molmet.2022.101605.

5. Covarrubias, A.J., Kale, A., Perrone, R., Lopez-Dominguez, J.A., Pisco, A.O., Kasler, H.G., Schmidt, M.S., Heckenbach, I., Kwok, R., Wiley, C.D., et al. (2020). Senescent cells promote tissue NAD(+) decline during ageing via the activation of CD38(+) macrophages. Nat Metab 2, 1265–1283. 10.1038/s42255-020-00305-3.

6. Gomes, A.P., Price, N.L., Ling, A.J., Moslehi, J.J., Montgomery, M.K., Rajman, L., White, J.P., Teodoro, J.S., Wrann, C.D., Hubbard, B.P., et al. (2013). Declining NAD(+) induces a pseudohypoxic state disrupting nuclear-mitochondrial communication during aging. Cell 155, 1624–1638. 10.1016/j.cell.2013.11.037.

7. Pirinen, E., Auranen, M., Khan, N.A., Brilhante, V., Urho, N., Pessia, A., Hakkarainen, A., Ulla Heinonen, J.K., Schmidt, M.S., Haimilahti, K., et al. (2020). Niacin Cures Systemic NAD(+) Deficiency and Improves Muscle Performance in Adult-Onset Mitochondrial Myopathy. Cell Metab 32, 144. 10.1016/j.cmet.2020.05.020.

8. Enzyme clustering accelerates processing of intermediates through metabolic channeling. (2014).

9. Changes in oxidative damage, inflammation and [NAD(H)] with age in cerebrospinal fluid. (2013).

10. The NAD+/sirtuin pathway modulates longevity through activation of mitochondrial UPR and FOXO signaling. (2013).

11. Frederick, D.W., Loro, E., Liu, L., Davila, A., Jr., Chellappa, K., Silverman, I.M., Quinn, W.J., 3rd, Gosai, S.J., Tichy, E.D., Davis, J.G., et al. (2016). Loss of NAD Homeostasis Leads to Progressive and Reversible Degeneration of Skeletal Muscle. Cell Metab 24, 269-282. 10.1016/j.cmet.2016.07.005.

12. In vivo NAD assay reveals the intracellular NAD contents and redox state in healthy human brain and their age dependences. (2015).

13. Hepatic NAD+ deficiency as a therapeutic target for non-alcoholic fatty liver disease in ageing. (2016).

14. Yoshino, J., Mills, K.F., Yoon, M.J., and Imai, S. (2011). Nicotinamide mononucleotide, a key NAD(+) intermediate, treats the pathophysiology of diet- and age-induced diabetes in mice. Cell Metab 14, 528–536. 10.1016/j.cmet.2011.08.014.

15. Specific ablation of Nampt in adult neural stem cells recapitulates their functional defects during aging. (2014).

16. NAD+ intermediates: The biology and therapeutic potential of NMN and NR. (2019).

17. Lopez-Otin, C., Blasco, M.A., Partridge, L., Serrano, M., and Kroemer, G. (2023). Hallmarks of aging: An expanding universe. Cell 186, 243–278. 10.1016/j.cell.2022.11.001.

18. Aguayo-Mazzucato, C. (2020). Functional changes in beta cells during ageing and senescence. Diabetologia 63, 2022–2029. 10.1007/s00125-020-05185-6.

19. Aguayo-Mazzucato, C., Andle, J., Lee, T.B., Jr., Midha, A., Talemal, L., Chipashvili, V., Hollister-Lock, J., van Deursen, J., Weir, G., and Bonner-Weir, S. (2019). Acceleration of beta Cell Aging Determines Diabetes and Senolysis Improves Disease Outcomes. Cell Metab 30, 129–142 e124. 10.1016/j.cmet.2019.05.006.

20. Bahour, N., Bleichmar, L., Abarca, C., Wilmann, E., Sanjines, S., and Aguayo-Mazzucato, C. (2023). Clearance of p16Ink4a-positive cells in a mouse transgenic model does not change beta-cell mass and has limited effects on their proliferative capacity. Aging (Albany NY) 15, 441–458. 10.18632/aging.204483.

21. Baker, D.J., Childs, B.G., Durik, M., Wijers, M.E., Sieben, C.J., Zhong, J., Saltness, R.A., Jeganathan, K.B., Verzosa, G.C., Pezeshki, A., et al. (2016). Naturally occurring p16(Ink4a)-positive cells shorten healthy lifespan. Nature 530, 184–189. 10.1038/nature16932.

22. Baker, D.J., Wijshake, T., Tchkonia, T., LeBrasseur, N.K., Childs, B.G., van de Sluis, B., Kirkland, J.L., and van Deursen, J.M. (2011). Clearance of p16Ink4a-positive senescent cells delays ageing-associated disorders. Nature 479, 232–236. 10.1038/nature10600.

23. Cha, J., Aguayo-Mazzucato, C., and Thompson, P.J. (2023). Pancreatic beta-cell senescence in diabetes: mechanisms, markers and therapies. Front Endocrinol (Lausanne) 14, 1212716. 10.3389/fendo.2023.1212716.

24. Miwa, S., Kashyap, S., Chini, E., and von Zglinicki, T. (2022). Mitochondrial dysfunction in cell senescence and aging. J Clin Invest 132. 10.1172/JCI158447.

25. Palmer, A.K., Gustafson, B., Kirkland, J.L., and Smith, U. (2019). Cellular senescence: at the nexus between ageing and diabetes. Diabetologia 62, 1835–1841. 10.1007/s00125-019-4934-x.

26. Palmer, A.K., Tchkonia, T., LeBrasseur, N.K., Chini, E.N., Xu, M., and Kirkland, J.L. (2015). Cellular Senescence in Type 2 Diabetes: A Therapeutic Opportunity. Diabetes 64, 2289–2298. 10.2337/db14-1820.

27. Thompson, P.J., Shah, A., Ntranos, V., Van Gool, F., Atkinson, M., and Bhushan, A. (2019). Targeted Elimination of Senescent Beta Cells Prevents Type 1 Diabetes. Cell Metab 29, 1045–1060 e1010. 10.1016/j.cmet.2019.01.021.

28. Wiley, C.D., and Campisi, J. (2021). The metabolic roots of senescence: mechanisms and opportunities for intervention. Nat Metab 3, 1290–1301. 10.1038/s42255-021-00483-8.

29. Wiley, C.D., Velarde, M.C., Lecot, P., Liu, S., Sarnoski, E.A., Freund, A., Shirakawa, K., Lim, H.W., Davis, S.S., Ramanathan, A., et al. (2016). Mitochondrial Dysfunction Induces Senescence with a Distinct Secretory Phenotype. Cell Metab 23, 303–314. 10.1016/j.cmet.2015.11.011.

30. Chini, C.C.S., Cordeiro, H.S., Tran, N.L.K., and Chini, E.N. (2024). NAD metabolism: Role in senescence regulation and aging. Aging Cell 23, e13920. 10.1111/acel.13920.

31. Hou, Y., Wei, Y., Lautrup, S., Yang, B., Wang, Y., Cordonnier, S., Mattson, M.P., Croteau, D.L., and Bohr, V.A. (2021). NAD(+) supplementation reduces neuroinflammation and cell senescence in a transgenic mouse model of Alzheimer’s disease via cGAS-STING. Proc Natl Acad Sci U S A 118. 10.1073/pnas.2011226118.

32. Verdin, E. (2015). NAD(+) in aging, metabolism, and neurodegeneration. Science 350, 1208–1213. 10.1126/science.aac4854.

33. Xie, N., Zhang, L., Gao, W., Huang, C., Huber, P.E., Zhou, X., Li, C., Shen, G., and Zou, B. (2020). NAD(+) metabolism: pathophysiologic mechanisms and therapeutic potential. Signal Transduct Target Ther 5, 227. 10.1038/s41392-020-00311-7.

34. Yoshino, M., Yoshino, J., Kayser, B.D., Patti, G.J., Franczyk, M.P., Mills, K.F., Sindelar, M., Pietka, T., Patterson, B.W., Imai, S.I., and Klein, S. (2021). Nicotinamide mononucleotide increases muscle insulin sensitivity in prediabetic women. Science 372, 1224–1229. 10.1126/science.abe9985.

35. Zhang, H., Ryu, D., Wu, Y., Gariani, K., Wang, X., Luan, P., D’Amico, D., Ropelle, E.R., Lutolf, M.P., Aebersold, R., et al. (2016). NAD(+) repletion improves mitochondrial and stem cell function and enhances life span in mice. Science 352, 1436–1443. 10.1126/science.aaf2693.

36. Kumari, R., and Jat, P. (2021). Mechanisms of Cellular Senescence: Cell Cycle Arrest and Senescence Associated Secretory Phenotype. Front Cell Dev Biol 9, 645593. 10.3389/fcell.2021.645593.

37. Campisi, J., Kapahi, P., Lithgow, G.J., Melov, S., Newman, J.C., and Verdin, E. (2019). From discoveries in ageing research to therapeutics for healthy ageing. Nature 571, 183–192. 10.1038/s41586-019-1365-2.

38. Sapieha, P., and Mallette, F.A. (2018). Cellular Senescence in Postmitotic Cells: Beyond Growth Arrest. Trends Cell Biol 28, 595–607. 10.1016/j.tcb.2018.03.003.

39. Aguayo-Mazzucato, C., van Haaren, M., Mruk, M., Lee, T.B., Jr., Crawford, C., Hollister-Lock, J., Sullivan, B.A., Johnson, J.W., Ebrahimi, A., Dreyfuss, J.M., et al. (2017). beta Cell Aging Markers Have Heterogeneous Distribution and Are Induced by Insulin Resistance. Cell Metab 25, 898–910 e895. 10.1016/j.cmet.2017.03.015.

40. Midha, A., Pan, H., Abarca, C., Andle, J., Carapeto, P., Bonner-Weir, S., and Aguayo-Mazzucato, C. (2021). Unique Human and Mouse beta-Cell Senescence-Associated Secretory Phenotype (SASP) Reveal Conserved Signaling Pathways and Heterogeneous Factors. Diabetes 70, 1098–1116. 10.2337/db20-0553.

41. Rubin de Celis, M.F., Garcia-Martin, R., Syed, I., Lee, J., Aguayo-Mazzucato, C., Bonner-Weir, S., and Kahn, B.B. (2022). PAHSAs reduce cellular senescence and protect pancreatic beta cells from metabolic stress through regulation of Mdm2/p53. Proc Natl Acad Sci U S A 119, e2206923119. 10.1073/pnas.2206923119.

42. Guarente, L., Sinclair, D.A., and Kroemer, G. (2024). Human trials exploring anti-aging medicines. Cell Metab 36, 354–376. 10.1016/j.cmet.2023.12.007.

43. Murao, N., Yokoi, N., Takahashi, H., Hayami, T., Minami, Y., and Seino, S. (2022). Increased glycolysis affects beta-cell function and identity in aging and diabetes. Mol Metab 55, 101414. 10.1016/j.molmet.2021.101414.

44. Metabolic regulation of transcription through compartmentalized NAD+ biosynthesis. (2018).

45. Lu, M.J., Busquets, J., Impedovo, V., Wilson, C.N., Chan, H.R., Chang, Y.T., Matsui, W., Tiziani, S., and Cambronne, X.A. (2024). SLC25A51 decouples the mitochondrial NAD(+)/NADH ratio to control proliferation of AML cells. Cell Metab 36, 808–821 e806. 10.1016/j.cmet.2024.01.013.

46. Kory, N., de Bos, J.U., van der Rijt, S., Jankovic, N., Gura, M., Arp, N., Pena, I.A., Prakash, G., Chan, S.H., Kunchok, T., et al. (2020). MCART1/SLC25A51 is required for mitochondrial NAD transport. Science Advances 6. ARTN eabe5310 10.1126/sciadv.abe5310.

47. Guldenpfennig, A., Hopp, A.K., Muskalla, L., Manetsch, P., Raith, F., Hellweg, L., Dordelmann, C., Leslie Pedrioli, D.M., Johnsson, K., Superti-Furga, G., and Hottiger, M.O. (2023). Absence of mitochondrial SLC25A51 enhances PARP1-dependent DNA repair by increasing nuclear NAD+ levels. Nucleic acids research. 10.1093/nar/gkad659.

48. Todisco, S., Agrimi, G., Castegna, A., and Palmieri, F. (2006). Identification of the mitochondrial NAD+ transporter in Saccharomyces cerevisiae. J Biol Chem 281, 1524–1531. 10.1074/jbc.M510425200.

49. Palmieri, F., Rieder, B., Ventrella, A., Blanco, E., Do, P.T., Nunes-Nesi, A., Trauth, A.U., Fiermonte, G., Tjaden, J., Agrimi, G., et al. (2009). Molecular Identification and Functional Characterization of Arabidopsis thaliana Mitochondrial and Chloroplastic NAD(+) Carrier Proteins. Journal of Biological Chemistry 284, 31249–31259. 10.1074/jbc.M109.041830.

50. Kory, N., Uit de Bos, J., van der Rijt, S., Jankovic, N., Gura, M., Arp, N., Pena, I.A., Prakash, G., Chan, S.H., Kunchok, T., et al. (2020). MCART1/SLC25A51 is required for mitochondrial NAD transport. Sci Adv 6. 10.1126/sciadv.abe5310.

51. Luongo, T.S., Eller, J.M., Lu, M.J., Niere, M., Raith, F., Perry, C., Bornstein, M.R., Oliphint, P., Wang, L., McReynolds, M.R., et al. (2020). SLC25A51 is a mammalian mitochondrial NAD(+) transporter. Nature 588, 174–179. 10.1038/s41586-020-2741-7.

52. Girardi, E., Agrimi, G., Goldmann, U., Fiume, G., Lindinger, S., Sedlyarov, V., Srndic, I., Gurtl, B., Agerer, B., Kartnig, F., et al. (2020). Epistasis-driven identification of SLC25A51 as a regulator of human mitochondrial NAD import. Nature Communications 11. ARTN 6145 10.1038/s41467-020-19871-x.

53. Orlandi, I., Stamerra, G., and Vai, M. (2018). Altered Expression of Mitochondrial NAD(+) Carriers Influences Yeast Chronological Lifespan by Modulating Cytosolic and Mitochondrial Metabolism. Front Genet 9, 676. 10.3389/fgene.2018.00676.

54. Casella, G., Munk, R., Kim, K.M., Piao, Y., De, S., Abdelmohsen, K., and Gorospe, M. (2019). Transcriptome signature of cellular senescence. Nucleic Acids Res 47, 7294–7305. 10.1093/nar/gkz555.

55. Marthandan, S., Priebe, S., Groth, M., Guthke, R., Platzer, M., Hemmerich, P., and Diekmann, S. (2015). Hormetic effect of rotenone in primary human fibroblasts. Immun Ageing 12, 11. 10.1186/s12979-015-0038-8.

56. Zirkel, A., Nikolic, M., Sofiadis, K., Mallm, J.P., Brackley, C.A., Gothe, H., Drechsel, O., Becker, C., Altmuller, J., Josipovic, N., et al. (2018). HMGB2 Loss upon Senescence Entry Disrupts Genomic Organization and Induces CTCF Clustering across Cell Types. Mol Cell 70, 730–744 e736. 10.1016/j.molcel.2018.03.030.

57. Santaguida, S., Richardson, A., Iyer, D.R., M’Saad, O., Zasadil, L., Knouse, K.A., Wong, Y.L., Rhind, N., Desai, A., and Amon, A. (2017). Chromosome Mis-segregation Generates Cell-Cycle-Arrested Cells with Complex Karyotypes that Are Eliminated by the Immune System. Dev Cell 41, 638–651 e635. 10.1016/j.devcel.2017.05.022.

58. Kovatcheva, M., Liao, W., Klein, M.E., Robine, N., Geiger, H., Crago, A.M., Dickson, M.A., Tap, W.D., Singer, S., and Koff, A. (2017). ATRX is a regulator of therapy induced senescence in human cells. Nat Commun 8, 386. 10.1038/s41467-017-00540-5.

59. Lammermann, I., Terlecki-Zaniewicz, L., Weinmullner, R., Schosserer, M., Dellago, H., de Matos Branco, A.D., Autheried, D., Sevcnikar, B., Kleissl, L., Berlin, I., et al. (2018). Blocking negative effects of senescence in human skin fibroblasts with a plant extract. NPJ Aging Mech Dis 4, 4. 10.1038/s41514-018-0023-5.

60. Helman, A., Avrahami, D., Klochendler, A., Glaser, B., Kaestner, K.H., Ben-Porath, I., and Dor, Y. (2016). Effects of ageing and senescence on pancreatic beta-cell function. Diabetes Obes Metab 18 *Suppl 1*, 58–62. 10.1111/dom.12719.

61. Steiner, D.J., Kim, A., Miller, K., and Hara, M. (2010). Pancreatic islet plasticity: interspecies comparison of islet architecture and composition. Islets 2, 135–145. 10.4161/isl.2.3.11815.

62. Tsuchitani, M., Sato, J., and Kokoshima, H. (2016). A comparison of the anatomical structure of the pancreas in experimental animals. J Toxicol Pathol 29, 147–154. 10.1293/tox.2016-0016.

63. Johmura, Y., Yamanaka, T., Omori, S., Wang, T.W., Sugiura, Y., Matsumoto, M., Suzuki, N., Kumamoto, S., Yamaguchi, K., Hatakeyama, S., et al. (2021). Senolysis by glutaminolysis inhibition ameliorates various age-associated disorders. Science 371, 265–270. 10.1126/science.abb5916.

64. Dai, C., Kayton, N.S., Shostak, A., Poffenberger, G., Cyphert, H.A., Aramandla, R., Thompson, C., Papagiannis, I.G., Emfinger, C., Shiota, M., et al. (2016). Stress-impaired transcription factor expression and insulin secretion in transplanted human islets. J Clin Invest 126, 1857–1870. 10.1172/JCI83657.

65. Hattangady, N.G., Carter, K., Maroni-Rana, B., Wang, T., Ayers, J.L., Yu, M., and Grady, W.M. (2024). Mapping the core senescence phenotype of primary human colon fibroblasts. Aging (Albany NY) 16, 3068–3087. 10.18632/aging.205577.

66. Lavandoski, P., Pierdona, V., Maurmann, R.M., Grun, L.K., Guma, F., and Barbe-Tuana, F.M. (2023). Eotaxin-1/CCL11 promotes cellular senescence in human-derived fibroblasts through pro-oxidant and pro-inflammatory pathways. Front Immunol 14, 1243537. 10.3389/fimmu.2023.1243537.

67. Zhang, Y., Tang, X., Wang, Z., Wang, L., Chen, Z., Qian, J.Y., Tian, Z., and Zhang, S.Y. (2023). The chemokine CCL17 is a novel therapeutic target for cardiovascular aging. Signal Transduct Target Ther 8, 157. 10.1038/s41392-023-01363-1.

68. Zhao, H., Liu, Z., Chen, H., Han, M., Zhang, M., Liu, K., Jin, H., Liu, X., Shi, M., Pu, W., et al. (2024). Identifying specific functional roles for senescence across cell types. Cell 187, 7314–7334 e7321. 10.1016/j.cell.2024.09.021.

69. Kawagoe, Y., Kawashima, I., Sato, Y., Okamoto, N., Matsubara, K., and Kawamura, K. (2020). CXCL5-CXCR2 signaling is a senescence-associated secretory phenotype in preimplantation embryos. Aging Cell 19, e13240. 10.1111/acel.13240.

70. Subramanian, A., Tamayo, P., Mootha, V.K., Mukherjee, S., Ebert, B.L., Gillette, M.A., Paulovich, A., Pomeroy, S.L., Golub, T.R., Lander, E.S., and Mesirov, J.P. (2005). Gene set enrichment analysis: a knowledge-based approach for interpreting genome-wide expression profiles. Proc Natl Acad Sci U S A 102, 15545–15550. 10.1073/pnas.0506580102.

71. Pullen, T.J., Khan, A.M., Barton, G., Butcher, S.A., Sun, G., and Rutter, G.A. (2010). Identification of genes selectively disallowed in the pancreatic islet. Islets 2, 89–95. 10.4161/isl.2.2.11025.

72. Thorrez, L., Laudadio, I., Van Deun, K., Quintens, R., Hendrickx, N., Granvik, M., Lemaire, K., Schraenen, A., Van Lommel, L., Lehnert, S., et al. (2011). Tissue-specific disallowance of housekeeping genes: the other face of cell differentiation. Genome Res 21, 95–105. 10.1101/gr.109173.110.

73. Miyazaki, J., Araki, K., Yamato, E., Ikegami, H., Asano, T., Shibasaki, Y., Oka, Y., and Yamamura, K. (1990). Establishment of a pancreatic beta cell line that retains glucose-inducible insulin secretion: special reference to expression of glucose transporter isoforms. Endocrinology 127, 126–132. 10.1210/endo-127-1-126.

74. Wang, Z., Wei, D., and Xiao, H. (2013). Methods of cellular senescence induction using oxidative stress. Methods Mol Biol 1048, 135–144. 10.1007/978-1-62703-556-9_11.

75. High passage MIN6 cells have impaired insulin secretion with impaired glucose and lipid oxidation. (2012).

76. Ni, R., Cao, T., Xiong, S., Ma, J., Fan, G.C., Lacefield, J.C., Lu, Y., Le Tissier, S., and Peng, T. (2016). Therapeutic inhibition of mitochondrial reactive oxygen species with mito-TEMPO reduces diabetic cardiomyopathy. Free Radic Biol Med 90, 12–23. 10.1016/j.freeradbiomed.2015.11.013.

77. Maechler, P., Jornot, L., and Wollheim, C.B. (1999). Hydrogen peroxide alters mitochondrial activation and insulin secretion in pancreatic beta cells. J Biol Chem 274, 27905–27913. 10.1074/jbc.274.39.27905.

78. Nacarelli, T., Lau, L., Fukumoto, T., Zundell, J., Fatkhutdinov, N., Wu, S., Aird, K.M., Iwasaki, O., Kossenkov, A.V., Schultz, D., et al. (2019). NAD(+) metabolism governs the proinflammatory senescence-associated secretome. Nat Cell Biol 21, 397–407. 10.1038/s41556-019-0287-4.

79. Prolla, T.A., and Denu, J.M. (2014). NAD+ deficiency in age-related mitochondrial dysfunction. Cell Metab 19, 178–180. 10.1016/j.cmet.2014.01.005.

80. Weiss-Sadan, T., Ge, M., Hayashi, M., Gohar, M., Yao, C.H., de Groot, A., Harry, S., Carlin, A., Fischer, H., Shi, L., et al. (2023). NRF2 activation induces NADH-reductive stress, providing a metabolic vulnerability in lung cancer. Cell Metab 35, 487–503 e487. 10.1016/j.cmet.2023.01.012.

81. Bryne, J.C., Valen, E., Tang, M.H., Marstrand, T., Winther, O., da Piedade, I., Krogh, A., Lenhard, B., and Sandelin, A. (2008). JASPAR, the open access database of transcription factor-binding profiles: new content and tools in the 2008 update. Nucleic Acids Res 36, D102–106. 10.1093/nar/gkm955.

82. Farre, D., Roset, R., Huerta, M., Adsuara, J.E., Rosello, L., Alba, M.M., and Messeguer, X. (2003). Identification of patterns in biological sequences at the ALGGEN server: PROMO and MALGEN. Nucleic Acids Res 31, 3651–3653. 10.1093/nar/gkg605.

83. Yamauchi, S., Sugiura, Y., Yamaguchi, J., Zhou, X., Takenaka, S., Odawara, T., Fukaya, S., Fujisawa, T., Naguro, I., Uchiyama, Y., et al. (2024). Mitochondrial fatty acid oxidation drives senescence. Sci Adv 10, eado5887. 10.1126/sciadv.ado5887.

84. Correia-Melo, C., Marques, F.D., Anderson, R., Hewitt, G., Hewitt, R., Cole, J., Carroll, B.M., Miwa, S., Birch, J., Merz, A., et al. (2016). Mitochondria are required for pro-ageing features of the senescent phenotype. EMBO J 35, 724–742. 10.15252/embj.201592862.

85. Goyal, S., Paspureddi, A., Lu, M.J., Chan, H.R., Lyons, S.N., Wilson, C.N., Niere, M., Ziegler, M., and Cambronne, X.A. (2023). Dynamics of SLC25A51 reveal preference for oxidized NAD(+) and substrate led transport. EMBO Rep 24, e56596. 10.15252/embr.202256596.

86. Guernsey, D.L., Jiang, H., Campagna, D.R., Evans, S.C., Ferguson, M., Kellogg, M.D., Lachance, M., Matsuoka, M., Nightingale, M., Rideout, A., et al. (2009). Mutations in mitochondrial carrier family gene SLC25A38 cause nonsyndromic autosomal recessive congenital sideroblastic anemia. Nat Genet 41, 651–653. 10.1038/ng.359.

87. Guldenpfennig, A., Hopp, A.K., Muskalla, L., Manetsch, P., Raith, F., Hellweg, L., Dordelmann, C., Leslie Pedrioli, D.M., Johnsson, K., Superti-Furga, G., and Hottiger, M.O. (2023). Absence of mitochondrial SLC25A51 enhances PARP1-dependent DNA repair by increasing nuclear NAD+ levels. Nucleic Acids Res 51, 9248–9265. 10.1093/nar/gkad659.

88. d’Adda di Fagagna, F., Reaper, P.M., Clay-Farrace, L., Fiegler, H., Carr, P., Von Zglinicki, T., Saretzki, G., Carter, N.P., and Jackson, S.P. (2003). A DNA damage checkpoint response in telomere-initiated senescence. Nature 426, 194–198. 10.1038/nature02118.

89. Anello, M., Lupi, R., Spampinato, D., Piro, S., Masini, M., Boggi, U., Del Prato, S., Rabuazzo, A.M., Purrello, F., and Marchetti, P. (2005). Functional and morphological alterations of mitochondria in pancreatic beta cells from type 2 diabetic patients. Diabetologia 48, 282–289. 10.1007/s00125-004-1627-9.

90. Aoyagi, K., Yamashita, S.I., Akimoto, Y., Nishiwaki, C., Nakamichi, Y., Udagawa, H., Abe, M., Sakimura, K., Kanki, T., and Ohara-Imaizumi, M. (2023). A new beta cell-specific mitophagy reporter mouse shows that metabolic stress leads to accumulation of dysfunctional mitochondria despite increased mitophagy. Diabetologia 66, 147–162. 10.1007/s00125-022-05800-8.

91. Eto, K., Tsubamoto, Y., Terauchi, Y., Sugiyama, T., Kishimoto, T., Takahashi, N., Yamauchi, N., Kubota, N., Murayama, S., Aizawa, T., et al. (1999). Role of NADH shuttle system in glucose-induced activation of mitochondrial metabolism and insulin secretion. Science 283, 981–985. 10.1126/science.283.5404.981.

92. Freeman, H.C., Hugill, A., Dear, N.T., Ashcroft, F.M., and Cox, R.D. (2006). Deletion of nicotinamide nucleotide transhydrogenase: a new quantitive trait locus accounting for glucose intolerance in C57BL/6J mice. Diabetes 55, 2153–2156. 10.2337/db06-0358.

93. Helman, A., Klochendler, A., Azazmeh, N., Gabai, Y., Horwitz, E., Anzi, S., Swisa, A., Condiotti, R., Granit, R.Z., Nevo, Y., et al. (2016). p16(Ink4a)-induced senescence of pancreatic beta cells enhances insulin secretion. Nat Med 22, 412–420. 10.1038/nm.4054.

94. Ishihara, H., Wang, H., Drewes, L.R., and Wollheim, C.B. (1999). Overexpression of monocarboxylate transporter and lactate dehydrogenase alters insulin secretory responses to pyruvate and lactate in beta cells. J Clin Invest 104, 1621–1629. 10.1172/JCI7515.

95. Kobiita, A., Silva, P.N., Schmid, M.W., and Stoffel, M. (2023). FoxM1 coordinates cell division, protein synthesis, and mitochondrial activity in a subset of beta cells during acute metabolic stress. Cell Rep 42, 112986. 10.1016/j.celrep.2023.112986.

96. Maassen, J.A., LM, T.H., Van Essen, E., Heine, R.J., Nijpels, G., Jahangir Tafrechi, R.S., Raap, A.K., Janssen, G.M., and Lemkes, H.H. (2004). Mitochondrial diabetes: molecular mechanisms and clinical presentation. Diabetes 53 *Suppl 1*, S103–109. 10.2337/diabetes.53.2007.s103.

97. Pinti, M.V., Fink, G.K., Hathaway, Q.A., Durr, A.J., Kunovac, A., and Hollander, J.M. (2019). Mitochondrial dysfunction in type 2 diabetes mellitus: an organ-based analysis. Am J Physiol Endocrinol Metab 316, E268–E285. 10.1152/ajpendo.00314.2018.

98. Rubi, B., del Arco, A., Bartley, C., Satrustegui, J., and Maechler, P. (2004). The malate-aspartate NADH shuttle member Aralar1 determines glucose metabolic fate, mitochondrial activity, and insulin secretion in beta cells. J Biol Chem 279, 55659–55666. 10.1074/jbc.M409303200.

99. Sakai, K., Matsumoto, K., Nishikawa, T., Suefuji, M., Nakamaru, K., Hirashima, Y., Kawashima, J., Shirotani, T., Ichinose, K., Brownlee, M., and Araki, E. (2003). Mitochondrial reactive oxygen species reduce insulin secretion by pancreatic beta-cells. Biochem Biophys Res Commun 300, 216–222. 10.1016/s0006-291x(02)02832-2.

100. Sekine, N., Cirulli, V., Regazzi, R., Brown, L.J., Gine, E., Tamarit-Rodriguez, J., Girotti, M., Marie, S., MacDonald, M.J., Wollheim, C.B., and, et al. (1994). Low lactate dehydrogenase and high mitochondrial glycerol phosphate dehydrogenase in pancreatic beta-cells. Potential role in nutrient sensing. J Biol Chem 269, 4895–4902.

101. Walker, E.M., Cha, J., Tong, X., Guo, M., Liu, J.H., Yu, S., Iacovazzo, D., Mauvais-Jarvis, F., Flanagan, S.E., Korbonits, M., et al. (2021). Sex-biased islet beta cell dysfunction is caused by the MODY MAFA S64F variant by inducing premature aging and senescence in males. Cell Rep 37, 109813. 10.1016/j.celrep.2021.109813.

102. Zhang, G.F., Jensen, M.V., Gray, S.M., El, K., Wang, Y., Lu, D., Becker, T.C., Campbell, J.E., and Newgard, C.B. (2021). Reductive TCA cycle metabolism fuels glutamine- and glucose-stimulated insulin secretion. Cell Metab 33, 804–817 e805. 10.1016/j.cmet.2020.11.020.

103. Krishnamurthy, J., Ramsey, M.R., Ligon, K.L., Torrice, C., Koh, A., Bonner-Weir, S., and Sharpless, N.E. (2006). p16INK4a induces an age-dependent decline in islet regenerative potential. Nature 443, 453–457. 10.1038/nature05092.

104. Segerstolpe, A., Palasantza, A., Eliasson, P., Andersson, E.M., Andreasson, A.C., Sun, X., Picelli, S., Sabirsh, A., Clausen, M., Bjursell, M.K., et al. (2016). Single-Cell Transcriptome Profiling of Human Pancreatic Islets in Health and Type 2 Diabetes. Cell Metab 24, 593–607. 10.1016/j.cmet.2016.08.020.

105. Qiu, W.L., Zhang, Y.W., Feng, Y., Li, L.C., Yang, L., and Xu, C.R. (2017). Deciphering Pancreatic Islet beta Cell and alpha Cell Maturation Pathways and Characteristic Features at the Single-Cell Level. Cell Metab 25, 1194–1205 e1194. 10.1016/j.cmet.2017.04.003.

106. Li, Y., Bie, J., Zhao, L., Song, C., Zhang, T., Li, M., Yang, C., and Luo, J. (2023). SLC25A51 promotes tumor growth through sustaining mitochondria acetylation homeostasis and proline biogenesis. Cell Death Differ 30, 1916–1930. 10.1038/s41418-023-01185-2.

107. Zhang, J., Tang, Y., Zhang, S., Xie, Z., Ma, W., Liu, S., Fang, Y., Zheng, S., Huang, C., Yan, G., et al. (2025). Mitochondrial NAD(+) deficiency in vascular smooth muscle impairs collagen III turnover to trigger thoracic and abdominal aortic aneurysm. Nat Cardiovasc Res. 10.1038/s44161-024-00606-w.

108. Kubben, N., and Misteli, T. (2017). Shared molecular and cellular mechanisms of premature ageing and ageing-associated diseases. Nat Rev Mol Cell Biol 18, 595–609. 10.1038/nrm.2017.68.

109. de Souza Chaves, I., Feitosa-Araujo, E., Florian, A., Medeiros, D.B., da Fonseca-Pereira, P., Charton, L., Heyneke, E., Apfata, J.A.C., Pires, M.V., Mettler-Altmann, T., et al. (2019). The mitochondrial NAD(+) transporter (NDT1) plays important roles in cellular NAD(+) homeostasis in Arabidopsis thaliana. Plant J 100, 487–504. 10.1111/tpj.14452.

110. Maechler, P. (2013). Mitochondrial function and insulin secretion. Mol Cell Endocrinol 379, 12–18. 10.1016/j.mce.2013.06.019.

111. Ramzy, A., Tuduri, E., Glavas, M.M., Baker, R.K., Mojibian, M., Fox, J.K., O’Dwyer, S.M., Dai, D., Hu, X., Denroche, H.C., et al. (2020). AAV8 Ins1-Cre can produce efficient beta-cell recombination but requires consideration of off-target effects. Sci Rep 10, 10518. 10.1038/s41598-020-67136-w.

112. Wang, Z., Zhu, T., Rehman, K.K., Bertera, S., Zhang, J., Chen, C., Papworth, G., Watkins, S., Trucco, M., Robbins, P.D., et al. (2006). Widespread and stable pancreatic gene transfer by adeno-associated virus vectors via different routes. Diabetes 55, 875–884. 10.2337/diabetes.55.04.06.db05-0927.

113. Xiao, X., Guo, P., Prasadan, K., Shiota, C., Peirish, L., Fischbach, S., Song, Z., Gaffar, I., Wiersch, J., El-Gohary, Y., et al. (2014). Pancreatic cell tracing, lineage tagging and targeted genetic manipulations in multiple cell types using pancreatic ductal infusion of adeno-associated viral vectors and/or cell-tagging dyes. Nat Protoc 9, 2719–2724. 10.1038/nprot.2014.183.

114. Jimenez, V., Ayuso, E., Mallol, C., Agudo, J., Casellas, A., Obach, M., Munoz, S., Salavert, A., and Bosch, F. (2011). In vivo genetic engineering of murine pancreatic beta cells mediated by single-stranded adeno-associated viral vectors of serotypes 6, 8 and 9. Diabetologia 54, 1075-1086. 10.1007/s00125-011-2070-3.

115. Kim, D.J., Chung, H., Ji, S.C., Lee, S., Yu, K.S., Jang, I.J., and Cho, J.Y. (2019). Ursodeoxycholic acid exerts hepatoprotective effects by regulating amino acid, flavonoid, and fatty acid metabolic pathways. Metabolomics 15, 30. 10.1007/s11306-019-1494-5.

116. Basu, S., Duren, W., Evans, C.R., Burant, C.F., Michailidis, G., and Karnovsky, A. (2017). Sparse network modeling and metscape-based visualization methods for the analysis of large-scale metabolomics data. Bioinformatics 33, 1545–1553. 10.1093/bioinformatics/btx012.

117. Chen, W.W., Freinkman, E., and Sabatini, D.M. (2017). Rapid immunopurification of mitochondria for metabolite profiling and absolute quantification of matrix metabolites. Nat Protoc 12, 2215–2231. 10.1038/nprot.2017.104.

118. Kong, B.S., Min, S.H., Lee, C., and Cho, Y.M. (2021). Mitochondrial-encoded MOTS-c prevents pancreatic islet destruction in autoimmune diabetes. Cell Rep 36, 109447. 10.1016/j.celrep.2021.109447.

119. Debacq-Chainiaux, F., Erusalimsky, J.D., Campisi, J., and Toussaint, O. (2009). Protocols to detect senescence-associated beta-galactosidase (SA-betagal) activity, a biomarker of senescent cells in culture and in vivo. Nat Protoc 4, 1798–1806. 10.1038/nprot.2009.191.

120. Ma, S., Sun, S., Geng, L., Song, M., Wang, W., Ye, Y., Ji, Q., Zou, Z., Wang, S., He, X., et al. (2020). Caloric Restriction Reprograms the Single-Cell Transcriptional Landscape of Rattus Norvegicus Aging. Cell 180, 984–1001 e1022. 10.1016/j.cell.2020.02.008.

121. Kanehisa, M., and Goto, S. (2000). KEGG: kyoto encyclopedia of genes and genomes. Nucleic Acids Res 28, 27–30. 10.1093/nar/28.1.27.

122. Butler, A., Hoffman, P., Smibert, P., Papalexi, E., and Satija, R. (2018). Integrating single-cell transcriptomic data across different conditions, technologies, and species. Nat Biotechnol 36, 411–420. 10.1038/nbt.4096.

123. Becht, E., McInnes, L., Healy, J., Dutertre, C.A., Kwok, I.W.H., Ng, L.G., Ginhoux, F., and Newell, E.W. (2018). Dimensionality reduction for visualizing single-cell data using UMAP. Nat Biotechnol. 10.1038/nbt.4314.

124. Aran, D., Looney, A.P., Liu, L., Wu, E., Fong, V., Hsu, A., Chak, S., Naikawadi, R.P., Wolters, P.J., Abate, A.R., et al. (2019). Reference-based analysis of lung single-cell sequencing reveals a transitional profibrotic macrophage. Nat Immunol 20, 163–172. 10.1038/s41590-018-0276-y.

125. Patil, A., and Patil, A. (2021). CellKb Immune: a manually curated database of hematopoietic marker gene sets from 7 species for rapid cell type identification. bioRxiv, 2020.2012.2001.389890. 10.1101/2020.12.01.389890.

126. Han, X., Wang, R., Zhou, Y., Fei, L., Sun, H., Lai, S., Saadatpour, A., Zhou, Z., Chen, H., Ye, F., et al. (2018). Mapping the Mouse Cell Atlas by Microwell-Seq. Cell 173, 1307. 10.1016/j.cell.2018.05.012.

127. Krentz, N.A.J., Lee, M.Y.Y., Xu, E.E., Sproul, S.L.J., Maslova, A., Sasaki, S., and Lynn, F.C. (2018). Single-Cell Transcriptome Profiling of Mouse and hESC-Derived Pancreatic Progenitors. Stem Cell Reports 11, 1551–1564. 10.1016/j.stemcr.2018.11.008.

128. Xu, M., Pirtskhalava, T., Farr, J.N., Weigand, B.M., Palmer, A.K., Weivoda, M.M., Inman, C.L., Ogrodnik, M.B., Hachfeld, C.M., Fraser, D.G., et al. (2018). Senolytics improve physical function and increase lifespan in old age. Nat Med 24, 1246–1256. 10.1038/s41591-018-0092-9.

129. Palmer, A.K., Xu, M., Zhu, Y., Pirtskhalava, T., Weivoda, M.M., Hachfeld, C.M., Prata, L.G., van Dijk, T.H., Verkade, E., Casaclang-Verzosa, G., et al. (2019). Targeting senescent cells alleviates obesity-induced metabolic dysfunction. Aging Cell 18, e12950. 10.1111/acel.12950.

130. Xin, Y., Kim, J., Okamoto, H., Ni, M., Wei, Y., Adler, C., Murphy, A.J., Yancopoulos, G.D., Lin, C., and Gromada, J. (2016). RNA Sequencing of Single Human Islet Cells Reveals Type 2 Diabetes Genes. Cell Metab 24, 608–615. 10.1016/j.cmet.2016.08.018.

131. Li, J., Klughammer, J., Farlik, M., Penz, T., Spittler, A., Barbieux, C., Berishvili, E., Bock, C., and Kubicek, S. (2016). Single-cell transcriptomes reveal characteristic features of human pancreatic islet cell types. EMBO Rep 17, 178–187. 10.15252/embr.201540946.

132. Muraro, M.J., Dharmadhikari, G., Grun, D., Groen, N., Dielen, T., Jansen, E., van Gurp, L., Engelse, M.A., Carlotti, F., de Koning, E.J., and van Oudenaarden, A. (2016). A Single-Cell Transcriptome Atlas of the Human Pancreas. Cell Syst 3, 385–394 e383. 10.1016/j.cels.2016.09.002.

133. Woo, H.R., Koo, H.J., Kim, J., Jeong, H., Yang, J.O., Lee, I.H., Jun, J.H., Choi, S.H., Park, S.J., Kang, B., et al. (2016). Programming of Plant Leaf Senescence with Temporal and Inter-Organellar Coordination of Transcriptome in Arabidopsis. Plant Physiol 171, 452–467. 10.1104/pp.15.01929.

134. Wang, J., Sang, Y., Jin, S., Wang, X., Azad, G.K., McCormick, M.A., Kennedy, B.K., Li, Q., Wang, J., Zhang, X., et al. (2022). Single-cell RNA-seq reveals early heterogeneity during aging in yeast. Aging Cell 21, e13712. 10.1111/acel.13712.

